# Headbutting goats self-inflict traumatic brain injury

**DOI:** 10.64898/2026.06.26.734585

**Authors:** AS Oyadeyi, C Smith, B Willeford, G Grissett-Hardwick, K Fizzano, WE Robinson, AG Sorace, A Osborne, S Samuel, I Campbell, A Srinivas, JE McConathy, J Bartels, S Lapi, NL Ackermans

## Abstract

Traumatic brain injury (TBI) is a characteristic feature of neurodegenerative diseases such as Alzheimer’s disease and chronic traumatic encephalopathy. Small animal models have been used to establish clinically relevant biomarkers of neuropathology, however, they show significant anatomical differences from humans and are affected by artificial experimental manipulations, making them often unsuitable for longitudinal study of repetitive mild TBI. There is a need for more diverse and translationally relevant animal models to better understand disease onset and development.

Building on a previous study of neuropathology in headbutting bovids in the wild, this pilot study investigated whether freely headbutting domestic goats, which naturally engage in low-intensity, high-frequency head impacts, accumulate measurable biomarkers of neurodegeneration in cerebrospinal fluid (CSF) and brain tissue. Over a six-month period, three male goats (*Capra hircus*) were allowed to freely headbutt under continuous video surveillance. Monthly CSF samples were collected, and concentrations of key neurodegeneration biomarkers were measured via multiplex immunoassays, including amyloid-β peptides (Aβ_40_, Aβ_42_), total and phosphorylated tau (tTau and pTau), glial fibrillary acidic protein (GFAP), S100 calcium-binding protein B (S100B), and neurofilament M (NF-M). Imaging studies were completed for non-invasive assessment of glucose metabolism ([^18^F]-FDG) and neuroinflammation via translocator protein (TSPO, [^18^F]DPA-714). Postmortem immunohistochemistry was conducted on prefrontal cortical tissues using antibodies targeting pTau, GFAP, and S100B. Head impact kinematics were quantified using horn-mounted accelerometer and inclinometer sensors that recorded linear acceleration, rotational velocity, and head orientation during naturally occurring headbutting events.

Several notable trends were observed. Phosphorylated tau as well as reactive astrocytes were detected in the brain tissue, mirrored by elevated GFAP detected in the CSF. PET TSPO did not illustrate a significant signal, however, PET FDG revealed frontal-dominant activity in all goats, and one with asymmetrical activation. Overall, the goats sustained 5,000-7,000 head impacts each over six months. Accelerometer measurements recorded forces up to 388 N and peak acceleration up to 16.5 g. This multi-modal observational study is the first to characterize neurodegeneration biomarkers and kinematics in headbutting goats. Even at one year old, the combination of pTau and gliosis in both the brain tissue and CSF indicates that the goat’s repetitive head impacts begin to show neurodegenerative consequences early in life. Likely, the severity of these consequences increases with headbutts and age, eventually resulting in chronic neurodegeneration. This system shows promise as a large-animal model for the longitudinal study of the onset and progression of neurodegenerative disease.

## INTRODUCTION

Neurodegeneration is a progressive and irreversible process characterized by the structural and functional deterioration of neurons (Joseph, 2025). Neurodegenerative diseases (NDDs) include traumatic brain injury (TBI), Alzheimer’s disease (AD), Parkinson’s disease, frontotemporal dementia, and chronic traumatic encephalopathy (CTE) (Wilson et al., 2023; Gadhave et al., 2024; Giannakis and Konitsiotis, 2025), and are major cause of illness and death worldwide. However, very few treatments exist to prevent, slow, or remedy these diseases (Wareham et al., 2022), and knowledge gaps remain regarding disease onset. A single head injury can almost double the risk of dementia as compared to background risk (Li et al., 2017; Fann et al., 2018; Schneider et al., 2021), making TBI a prominent risk factor for NDDs, with at least 3% of dementia cases attributable to TBI (Livingston et al., 2020; Maas et al., 2022) and evidence of higher risk amongst contact-sport athletes. For example, at least 17% of repetitive TBI cases are reported to develop NDDs like CTE in boxers (Stiller and Weinberger, 1985), although in reality, the rate for athletes with repetitive TBI is likely much higher, as it has not been explicitly measured (McKee et al., 2009).

Each NDD exhibits distinct pathological features, severity, and progression rate but share some molecular mechanisms, which can offer targeted avenues for research. These convergences include the accumulation of misfolded proteins, leading to the formation of insoluble aggregates such as extracellular amyloid-beta (Aβ) plaques, intracellular neurofibrillary tangles composed of hyperphosphorylated tau, and Lewy bodies composed of aggregated alpha-synuclein (Soto, 2003; Wareham et al., 2022; Wilson et al., 2023). These protein aggregates disrupt cellular homeostasis, impair synaptic function, and can induce misfolding and aggregation of other proteins in a chain reaction, similarly to how pathological prion proteins aggregate (Frost and Diamond, 2010), thereby exacerbating neuronal damage (Sweeney et al., 2017). Ultimately, neurodegeneration is often the result of a combination of energy failure, oxidative stress, neuroinflammation, failure of proteostasis, genetic mutations, and glutamate excitotoxicity (Kim et al., 2015; Ghosh and Saadat, 2023).

Currently, state-of-the-art NDD diagnosis relies on a combination of fluid and tissue-based biomarkers, alongside neurobehavioral assessments (Anoop et al., 2010; Ahmad et al., 2023; Agnello et al., 2024; Beard et al., 2024). Imaging approaches have emerged to characterize NDDs, including positron emission tomography (PET) molecular imaging for associated biomarkers and magnetic resonance imaging (MRI) to assess disease-related anatomical alterations. Immunohistochemistry is used to confirm diagnosis postmortem by mapping the pathological extent of disease (Kleinman et al., 1994; Hatanpaa et al., 2008; Dugger and Dickson, 2017). However, diagnosis *in vivo* is difficult and NDD biomarkers are needed to improve disease detection and intervention, especially at early stages (DeKosky and Marek, 2003).

### Biomarkers for neurodegeneration

Tau is protein that contributes to the structural rigidity of the microtubules in axons (Avila et al., 2004). After a TBI, resulting neuronal damage causes intracellular release tau, and both total tau (tTau) and phosphorylated tau (pTau), are detected in elevated levels in cerebrospinal fluid (CSF) (Rubenstein et al., 2024), blood (Rubenstein et al., 2017), saliva (Olczak et al., 2019), and brain tissue, even for years after a single TBI (Johnson et al., 2012). Amyloid beta 42 (Aβ_42_) can also be detected at decreased levels after TBI and may correlate with clinical outcomes (Bogoslovsky et al., 2016). Therefore, tau and Aβ are used as biomarkers in both fluids and tissue, especially in association with monitoring NDD development. Histologically, the appearance of pTau in neurites, neuritic threads, neuritic plaques, pretangles, neurofibrillary tangles, glia, and other phosphorylated tau-rich conformations can be indicative of NDDs (Ferrer et al., 2014; Moloney et al., 2021). The specific distribution of these pTau conformations determines whether the tau pathology is primarily age related (PART) or related to rmTBI. The physical forces resulting from head trauma concentrate at the base of the sulci, resulting in a diagnostic pattern of cortical neuronal degradation, and therefore neuronal pTau accumulation, in the sulcal depths and perivascularly (McKee et al., 2023). Aside from tau inclusions, TBIs also incur glial abnormalities such as proliferation and hypertrophy especially within 24 hours post-injury (Mira et al., 2021). Additionally, tau and glia pathology often occur in combination in repetitive TBI pathology (Cherry et al., 2016).

It is currently unknown how severe or frequent TBI must be to cause long-term effects including development of NDDs like AD and CTE. AD is particularly difficulty to prevent, detect, and therefore treat, as current diagnostic techniques such as fluid biomarkers and PET imaging allow for detection at the earliest about a decade before severe symptoms appear whereas MRI diagnosis only appears alongside behavioral change in the late stages of disease (Jack, 2022). Genetic predisposing factors aside, due to the current limited resolution of diagnostic techniques and their invasiveness, early AD onset is not yet possible to detect. Therefore, the current focus for preventing, reducing, and treating these diseases is on acquiring a better understanding disease onset and development.

The complexity of in vivo measurements in humans combined with the lack of model translation has led to gaps in our knowledge specifically in the timeframe between acquiring a TBI and developing a clinically-diagnosable NDD. For example, for clinical diagnosis, the pathological profile of important neurodegeneration biomarkers of rmTBI is unclear due to moderate-severe human TBI cases being presented at the hospital while mTBI cases that lack overt clinical issues and may not seek treatment (Rahim et al., 2022). Moreover, most TBI biomarkers are used for research purposes only (Maas et al., 2022). Complicating this issue is the absence of well-defined reference values for neurodegeneration biomarkers (Schindler et al., 2024a), as biomarker concentrations vary depending on the identity of the biomarker, body fluid source, patient population, severity of injury, time of sample collection, assay method, and possibly other factors (Gardner et al., 2022; Ghaith et al., 2022; Rahim et al., 2022; Whitehouse et al., 2022). In addition, there are only few controlled studies examining longitudinal temporal changes of these biomarkers (Schnakers et al., 2021) in both human cohorts and animal models, and in these models standardized species-specific reference values are lacking (Giangrande et al., 2023).

### Difficulties in common laboratory model translation

Despite rats and mice being the most extensively used laboratory model species due to their genetic manipulability, well-characterized neuroanatomy, and cost-effectiveness (Dawson et al., 2018), NDD-associated proteinopathies do not naturally occur in these rodent models. Rodents are unable to develop the full spectrum of human-like neuropathological features even after repeated or severe injury, due to shorter lifespans and because the natural aging process in rodents does not involve development of amyloid plaques (Poon et al., 2020). Therefore, several experimental paradigms have been developed to model AD & TBI in rodent models. Transgenic rodent models can use humanized Aβ strains to model AD (Sasaguri et al., 2017) and weight drop, fluid percussion, or blast injury combined with transgenic models are used to model TBI in rodents (Xiong et al., 2013).

Despite the advances these models have brought to the field, clinical translation from rodent models to human cures has been challenging due to a myriad of species differences that include anatomy, physiology, injury parameters, and endpoints assessed (Ransohoff, 2018; Lisi et al., 2023). Brain size, gyrification, white-to-gray matter ratio, and neurovascular organization are highly variable between models, yet crucial in understanding the biomechanical force distribution and resulting neuropathological pattern caused by varying levels of TBI. Additionally, rotational acceleration is a primary driver of human brain injury, but is difficult to replicate at comparable levels due to the smaller mass and rigidity in small animal models (Rowson et al., 2012). Importantly, between 70-90% of human TBIs are mild and repetitive (Bielanin et al., 2024), occurring over a period of about 15 years on average (Stein et al., 2015). A review of long-term TBI survival studies indicated very few studies include outcome assessments beyond one month, with yet fewer extending to one year, of those, 96% were performed in rat and mouse models (Osier et al., 2015). A narrow timeframe of data collection caused by experimental restrictions, has limited our understanding of long-term consequences. Indeed, one meta-analysis indicated that in humans, TBI-related cognitive impairments can continue 1 - 4.5 years postinjury (Ruttan et al., 2008) and that mild repetitive head impacts (RHIs) have been linked to chronic NDDs such as AD, PD, and CTE (Bailes et al., 2014; McKee et al., 2016; van der Staay et al., 2017; Vink, 2018; Brett et al., 2022).

### A case for increased animal model diversity

TBI does not occur solely in humans (Tobiansky et al., 2026). Neuroscientists have clearly identified a need for increased species diversification in animal models (Brenowitz and Zakon, 2015; Hale, 2019) as a tool for improving translational research and addressing translational gaps (Garner, 2014; Lambert, 2023), and the NIH has stated that diversification of model species is an important milestone for increasing the predictive validity of AD models (National Institute of Aging, 2018). To expand model diversity, one can consider studying diverse model species in the lab, but also acquiring a better understanding of brain injuries in a natural context.

While no model can completely replicate the complexity of human neurodegeneration (Morganti-Kossmann et al., 2010), research on non-traditional model species has expanded our knowledge. For example, a gyrencephalic ferret model has shown histopathological and behavioral abnormalities following TBI (Schwerin et al., 2017; Schwerin et al., 2021). Histopathological markers of neurodegeneration that do not naturally occur in rodents are known to develop with age or injury in many vertebrate species, including primates, carnivores, cetartiodactyls, perissodactyls, rodents, and birds (Youssef et al., 2016; De Sousa et al., 2023; Ferrer, 2024; Ford et al., 2025). Larger model animals such as sheep and goats are known to naturally develop amyloid plaques and phosphorylated tau tangles (Braak et al., 1994; Nelson et al., 1994; Nelson and Saper, 1995; Reid et al., 2017; Davies et al., 2022; Nakayama et al., 2025), and have also been used as experimental models for NDDs with experimentally-induced TBI (Lewis et al., 1996; Finnie, 1997; Anderson et al., 1999; Anderson et al., 2003; Li et al., 2014). Sheep have been validated for cognition studies and may offer avenues towards better understanding cognitive decline associated with NDDs (Morton and Avanzo, 2011). As compared to rodents, the goat’s gyrencephalic brain is more comparable to the human brain in size and conformation (Schmidt et al., 2012), with moderate development of the nervous system relative to size (Roth and Dicke, 2005), and a significant white matter proportion of brain (25–30%) making them likely to exhibit axonal stretch and shear injury patterns that resemble those seen in human rmTBI, particularly when subjected to rotational forces. In addition, goats have a well-developed corpus callosum that enables cross-species comparisons of axonal pathology, including biomarker leakage and histopathological assessments of cytoskeletal integrity (Cernak, 2005). Additionally, goats can accumulate many of the biomarkers commonly used to detect acute or chronic TBI in humans, as markers for tau, beta amyloid, TDP-43, S100B, NSE, GFAP, Iba1, NeuN, Collagen IV, MAP2 and more have been detected in caprid brain tissue, CSF, and plasma (Tan et al., 2024). While goats and sheep are not identical species, they share similarities that allow findings in one species to inform the other.

### Previous neurodegeneration studies on Caprinae

Only six studies outside of our group have investigated the status of phosphorylated tau in brain tissue non-experimental Caprinae (subfamily of medium-sized bovids including goats and sheep) using immunohistochemistry. In two studies, no pTau was detected in goat, sheep, or serow < 4 yr (Nelson et al., 1994; Nakayama et al., 2025), however, Davies et al. (2022) detected pTau in female sheep in the same age range at 1 yr (Table 1). Previous CSF biomarker studies in sheep found the CSF Aβ_42/40_ ratio and tau levels to be comparable to human samples (Reid et al., 2017) and Aβ_42/40_ genes are identical between humans, sheep, and goats, with all of the main APP fragments found in humans detectable in goat CSF (Reid et al., 2017) (Table 2).

**Table 1.**
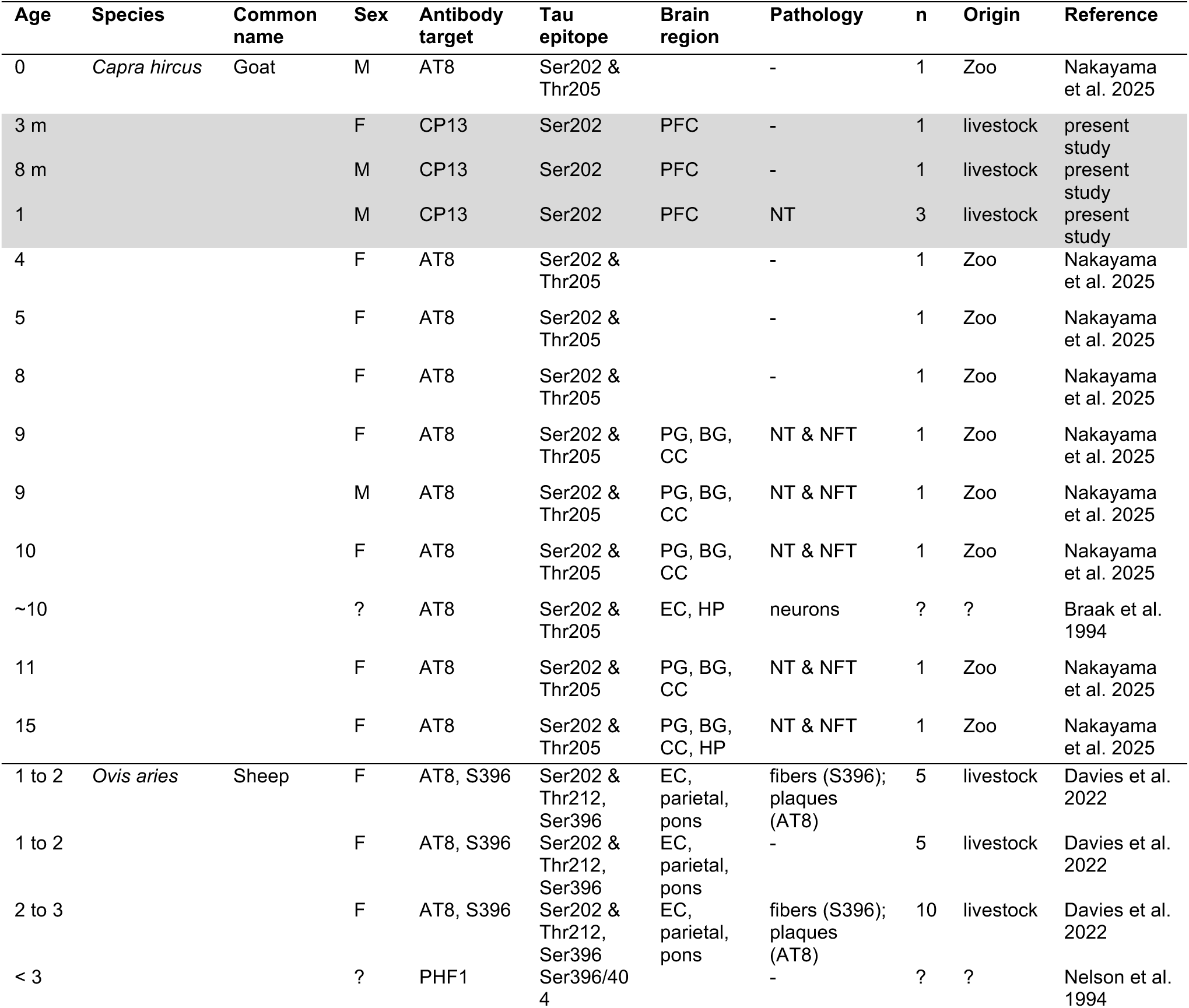
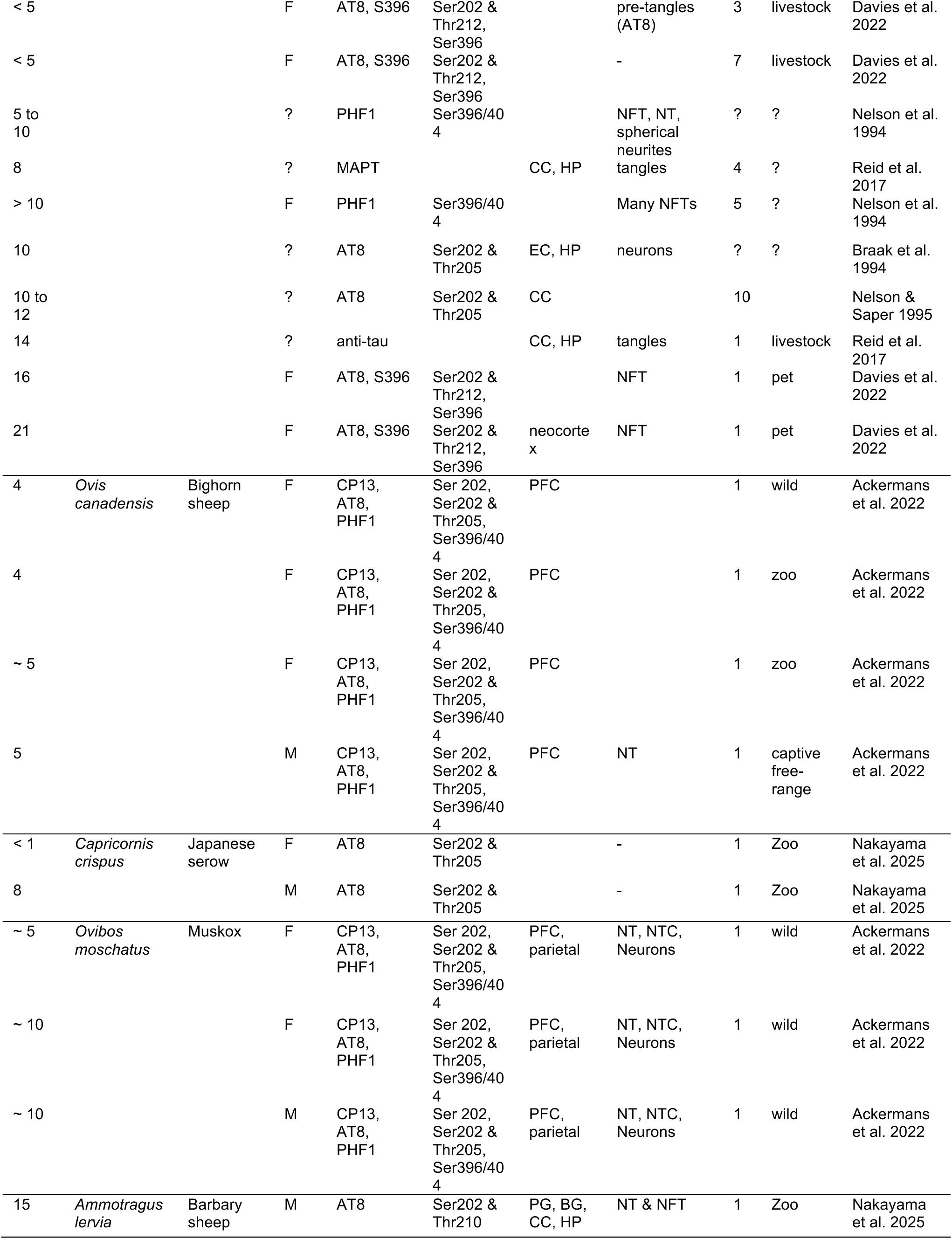
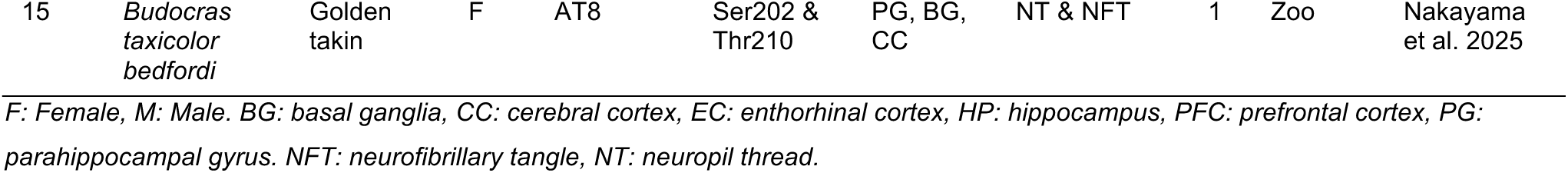
Summary of previous research investigating tau in Caprinae. Results from the present study are highlighted in grey.

**Table 2.** Percentage identity between key human neurodegeneration-related protein sequence and the goat reference sequence, indicating high homology. BLAST analysis used goat (*Capra hircus*) taxid 9925.

| Gene name | Goat vs. human % identity | Human sequence (length) | Goat sequence (length) | Coverage % |
| --- | --- | --- | --- | --- |
| APP770 | 97.01% | NP_000475.1 (770) | XP_017906012.1 (770) | 100 |
| Aβ1–40 | 100% | NP_000475.1 (770) | XP_017906012.1 (770) | 100 |
| Aβ1–42 | 100% | NP_000475.1 (770) | XP_017906012.1 (770) | 100 |
| α-cleavage site | 100% | NP_000475.1 (770) | XP_017906012.1 (770) | 100 |
| β-cleavage site | 100% | NP_000475.1 (770) | XP_017906012.1 (770) | 100 |
| γ-cleavage site | 100% | NP_000475.1 (770) | XP_017906012.1 (770) | 100 |
| BACE1 | 98.40% | NP_036236.1 (501) | XP_013824921.2 (501) | 100 |
| PSEN1 | 91.03% | NP_000012.1 (467) | XP_005686102.1 (468) | 100 |
| PSEN2 | 97.33% | NP_000438.2 (448) | XP_005690593.1 (449) | 100 |
| APOE | 70.35% | NP_000032.1 (317) | XP_017918050.1 (316) | 100 |
| APOE4 (112 position) | 100% | NP_000032.1 (317) | XP_017918050.1 (316) | 100 |
| APOE4 (158 position) | 100% | NP_000032.1 (317) | XP_017918050.1 (316) | 100 |
| NF-M | 81.46% | NP_005373.2 (916) | XP_013821584.2 (922) | 100 |
| MAPT (tau isoform 1) | 86.59% | NP_058519.3 (758) | XP_017920770.1 (601) | 81 |
| Tau isoform 1N3R | 90.34% | NP_001091180.1 (381) | XP_017920787.1 (372) | 100 |
| Tau isoform 2N3R | 91.06% | NP_001116539.1 (412) | XP_017920784.1 (403) | 100 |
| Tau isoform 1N4R | 90.11% | NP_001346197.1 (352) | XP_017920788.1 (343) | 100 |
| Tau isoform 2N4R | 90.52% | NP_005901.2 (441) | XP_017920782.1 (432) | 100 |
| GFAP | 95.14% | NP_002046.1 (432) | XP_017920743.1 (428) | 100 |
| S100B | 98.91% | NP_006263.1 (92) | XP_005675734.1 (92) | 100 |

All six tau isoforms are expressed in the sheep brain and can become hyperphosphorylated and involved in tau pathology. MAPT, however, differs between species, as compared to humans, goats lack exons 3, 6, and 8 (Nelson et al., 1996; Janke et al., 1999). Both 3R & 4R human tau isoforms are similarly expressed in sheep and goats, as opposed to mice, which only express 4R tau (Davies et al., 2022). Thus, sheep and goat models share many similarities with humans, making them relevant to the application of many methods used for clinical molecular NDD disease detection.

In addition to sharing anatomical and molecular similarities with humans, sheep and goat models offer a crucial advantage over any other model: the potential to induce their own brain damage through naturally occurring headbutting behavior. Our group was the first to discover chronic TBI pathology in brain tissue of wild bighorn sheep and muskoxen (Ackermans et al., 2022), species that had previously been assumed to be entirely resistant to TBI caused by repeated headbutting (Eck, 2014; Drake et al., 2016; Ackermans, 2025). The pathology severity in these species indicates that other headbutting bovids such as domestic goats, likely also accumulate neurodegeneration from repeated head impact behavior. Therefore, the present multimodal observational study investigated whether the goat’s natural headbutting behavior causes brain damage by measuring longitudinal clinical markers of disease *in vivo* and confirming with postmortem diagnosis.

Studying rmTBI acquired via headbutting may be the closest manner at present to replicate the development of complex pathology caused by TBI in real-world injury. Importantly, the headbutting goat model provides the opportunity to address two important research gaps by allowing for investigation of early stages of disease and long-term disease development form young to aged individuals. This is because headbutting avoids the artefacts traditionally associated with experimentally induced TBI such as surgery, anesthesia, and short-term endpoints.

Overall, this study represents the first report and attempt to quantitatively assess clinical neurodegeneration biomarkers in a population of headbutting domestic goats, including the first PET/MRI on goat brain. Our goal was to better understand neurodegenerative disease onset and set a baseline for future work in this model.

## METHODOLOGY

### Study design

The current study was designed as an observational cohort study in which individuals with different levels of exposure to headbutting were followed longitudinally to determine the incidence of brain damage as a dose-response measure to repeated headbutting over time. This type of design allows for the study of several outcomes within the same study (Jepsen et al., 2004). As this was a pilot study, animal numbers were limited. Funding source, animals, and imaging were dispersed across three separate institutions and two states. Involved personnel met virtually to organize the complex project. Both AL and MS stare veterinarians were consulted, as USDA-covered species crossed state lines. IACUCs at all institutions were consulted and approved all protocols. Throughout the process, each group was assigned specific roles and used open, clear communication. The authors recommend this type of model for the success of a multi-institutional project.

### Animals

Three male domestic goats (*Capra hircus*, n = 3), hereafter referred to as Goat A, Goat B, and Goat C, were acquired from local breeders and housed at Mississippi State University (MSU), in an indoor, climate-controlled enclosure with bedding and access to fresh water, hay, and feed. Animals were aged by teeth and estimated to be approximately six months old at the start of the experiment, and therefore approximately 1 yr at the end of the experiment. This study was approved by the MSU Institutional Animal Care and Use Committee (IACUC protocol #23-243). Animal husbandry was provided by veterinary staff and technicians within the AAALAC-accredited facility at MSU. Before the start of experimental procedures, the animals underwent a 14-day acclimatization period on arrival, where they were separated physically to prevent headbutting, but not visually, and they were monitored for any signs of illness. They were subsequently brought together in a pen and allowed to freely interact for seven months under continuous video recording. The goats underwent a baseline CT scan at the beginning of the experiment, followed by a PET/MRI brain scans at both the beginning with [^18^F]DPA-714 (6 mo) and end of the experiment with [^18^F]FDG (1 yr old) under anesthesia. The animals also underwent monthly sample collections of saliva, blood, and CSF for biomarker analysis. Animals were sedated prior to CSF collection only. After the final PET/MRI scan acquisition, the animals were humanely euthanized (Euthasol®, Vibrac 1 ml/4.5 kg) (Fig. 1).

**Figure 1.**
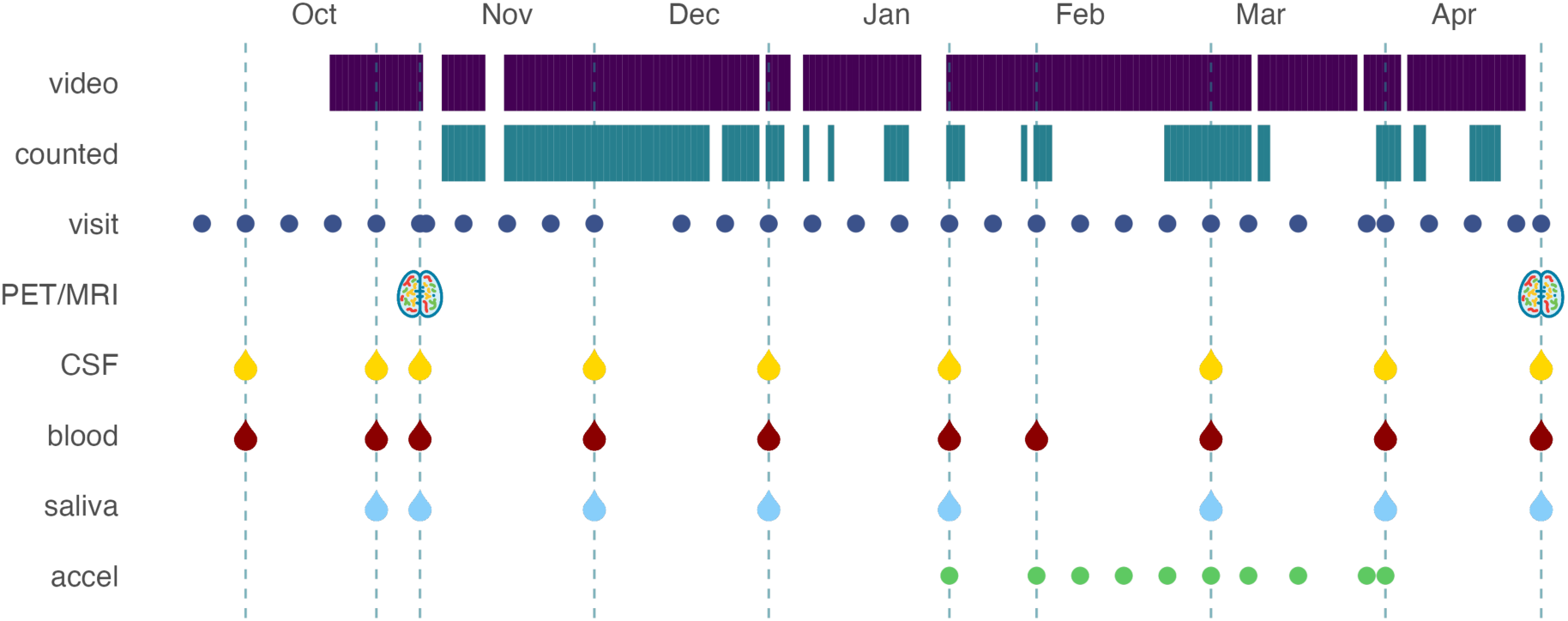
Timeline of a longitudinal study on headbutting goats, highlighting the different data modalities and collection timepoints. All three goats followed the same experimental schedule over seven months. Accel = accelerometer data collection.

An autopsy saw (Mopec 1000, with BD101 blade) and a chisel were used to remove the skull in the occipital region, vertically between the horns and the foramen magnum, and horizontally between the caudal aspect of the zygomatic processes of the temporal bone. The meninges, blood vessels and nerves were manually severed, and the brain was carefully extracted. Following best practice guidelines (Hyman et al., 2012; National Cancer Institute, 2016), the brain’s right hemisphere was transported on dry ice and then flash-frozen in liquid nitrogen before being stored at -80 ^°^C. The left hemisphere was fixed in 10 % paraformaldehyde at a 1:10 ratio. Fixed brain was blocked in preparation for immunohistochemistry (IHC) and sections were cut from a prefrontal cortex block on a vibratome (Leica S1000) at 50 μm thickness and stored in phosphate-buffered saline (PBS, pH 7.0) containing 0.1% sodium azide for preservation.

Two additional young goat brains were obtained from MSU as immunohistochemistry comparison specimens. One was a 3-month-old female, and the other was an 8-month-old male. Both goats were euthanized outside of this study as part of routine veterinary practice and originate from livestock herds.

### Immunohistochemistry

Some of the antibodies used in this study were designed to target human antigens, therefore their reactivity with goat sample tissues was confirmed through The Basic Local Alignment Search Tool (BLAST) NCBI database (Altschul et al., 1990). We compared DNA for non-redundant proteins against the *Capra hircus* taxid 9925 and a ranking of the different isoforms was performed based on sequence similarity and molecular weight, with percent identity and E-values is provided in Table 2. Homology values below 85% were considered low, above 85% moderate, and above 95% acceptable.

Sections from the anterior prefrontal cortex of the right hemispheres of each goat were stained using free-floating immunohistochemistry (IHC), building on the technique established in Ackermans et al. (2022), restated here with some adaptations.

We used antibodies raised against pS202 (CP13) to detect early pTau tangles and against pS396/404 (PHF-1) to detect late pTau tangle developments (Ghatamaneni et al., 2025). To identify astrocytes at different stages of reactivity, we used antibodies raised against GFAP to detect reactive astrocytes and S100B for broader mature astrocyte labeling (see Table 3 for full antibody details) (Steiner et al., 2007). We used primary antibody controls (sections where the primary antibody was not introduced) to rule out non-specific binding by the secondary antibodies.

**Table 3.**
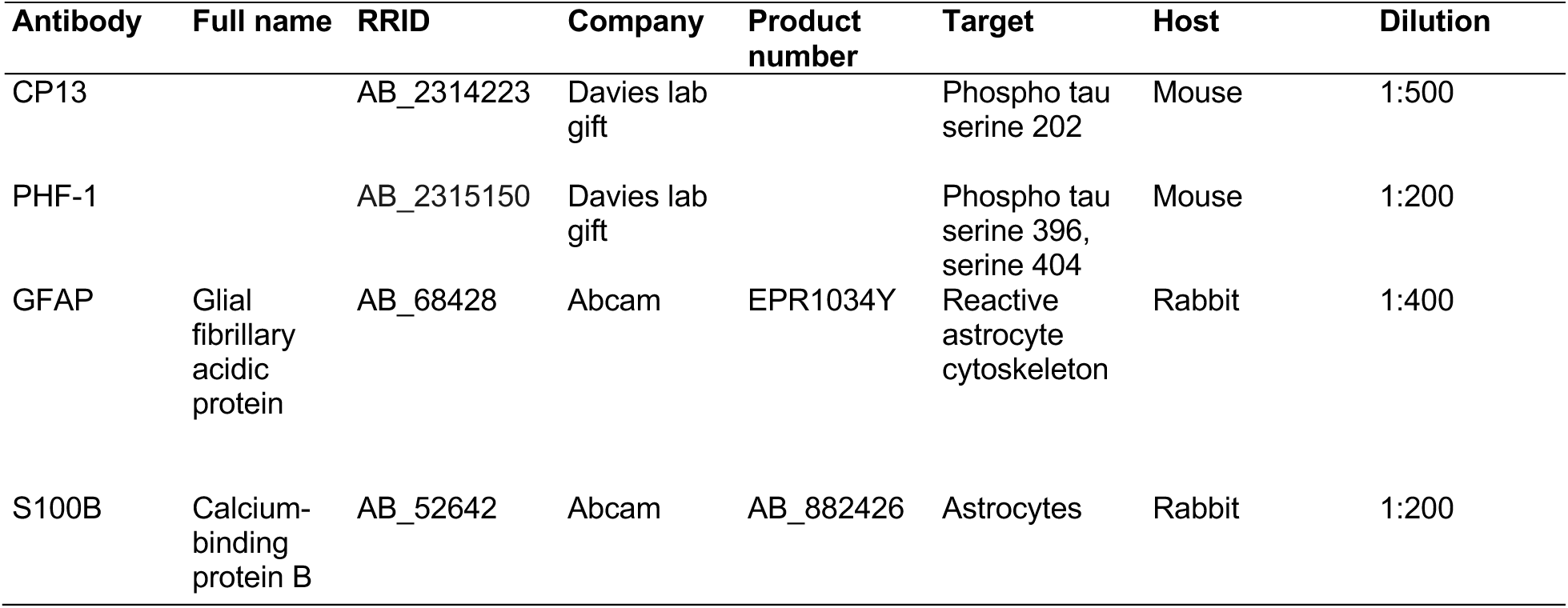
List of antibodies used in this study.

| Antibody | Full name | RRID | Company | Product number | Target | Host | Dilution |
| --- | --- | --- | --- | --- | --- | --- | --- |
| CP13 |  | AB_2314223 | Davies lab gift |  | Phospho tau serine 202 | Mouse | 1:500 |
| PHF-1 |  | AB_2315150 | Davies lab gift |  | Phospho tau serine 396, serine 404 | Mouse | 1:200 |
| GFAP | Glial fibrillary acidic protein | AB_68428 | Abcam | EPR1034Y | Reactive astrocyte cytoskeleton | Rabbit | 1:400 |
| S100B | Calcium-binding protein B | AB_52642 | Abcam | AB_882426 | Astrocytes | Rabbit | 1:200 |

Prior to antigen retrieval, sections were washed in phosphate-buffered saline (PBS), or Tris-buffered saline (TBS, pH 7.0) in the case of the CP13 protocol, four times for 5 minutes per wash on a shaker at 80 rpm. For the GFAP protocol, heat-mediated antigen retrieval was performed by suspending the slices in 10 mM citrate buffer (pH 8.75) in a water bath maintained at 85 °C for 30 minutes. This was followed by a 30-minute cool-down in the same buffer at room temperature. The sections were then transferred to a 24-well plate dish and washed four times as previously described. Sections were then incubated in 0.3% hydrogen peroxide with 0.3% Triton X-100 for 30 minutes to block endogenous peroxidase activity. After another wash step, non-specific binding was blocked by incubating the sections in 5% normal donkey serum (JacksonImmunoresearch, AB7475) for one hour. Incubation with primary antibody was performed overnight at 4 °C with the primary antibodies diluted in buffer containing 5% serum and 0.3% Triton X-100 and simultaneously, primary antibody control sections were processed using buffers without primary antibodies. After washing off primary antibodies, sections were incubated in biotinylated secondary antibodies, either donkey anti-mouse (Invitrogen, PA1-28627) or donkey anti-rabbit (JacksonImmunoresearch, AB_2340593) diluted 1:1000 in buffers with 5% serum and 0.3% Triton X-100 for one hour at room temperature. After secondary antibody incubation, sections were incubated with an avidin-biotin complex (Vectastain ABC kit, Vector Laboratories) and visualized using DAB peroxidase substrate (Vector laboratories, SK-4100) according to the manufacturer’s instructions. Sections were rinsed once in PBS or TBS and mounted on poly-L-Lysine-coated slides that were set to dry overnight. Finally, sections were counterstained with cresyl violet, dehydrated through an ethanol gradient, and cover slips were mounted with DPX mounting medium (Electron microscopy sciences).

### Microscopy and quantification

Microscopy images were taken on a Zeiss Axio Imager 2 brightfield microscope (Carl Zeiss Microscopy GMBG, Jena, Germany), with a 2.5x/0,085 objective (Zeiss EC Plan-Neofluar) and a 40x/1,3 oil objective (Zeiss EC Plan-Neofluar) using StereoInvestigator (version 2023.2.1, MBF Bioscience, Williston, VT, USA).

Semi-quantitative assessment of structures expressing pTau was performed using the optical fractionator workflow in StereoInvestigator (Counting frame size 230 µm x 230 µm, SRS grid layout 100%, Guard zone: 2 µm, optical dissector height: 30 µm), on each goat, with 1-4 non-consecutive sections per specimen from the prefrontal cortex, each counted exhaustively. Markers were placed digitally for neuropil threads expressing pTau, defined as axonal or dendric filaments made of abnormally-phosphorylated tau protein. Marker coordinates were exported to create density heatmaps for each section using R studio (Posit team, 2025).

To quantify sulcal patterns in relation to structures expressing pTau, sulcal depths were delimitated as the bottom 1/3 of the sulcus and tau-immunoreactivity was quantified in each region (1/3 and remaining 2/3 of sulcus) as in Hsu et al. (2018) and (Ackermans et al., 2022), using StereoInvestigator and the same program settings above. Similarly, to quantify association of tau-immunoreactive structures to blood vessels, vessels > 30 μm in diameter were counted exhaustively at 2.5 x magnification and the percentage of structures expressing pTau within 100 μm of the edge of the vessel was quantified and their distance from the vessel was reported.

### Biomarker sample collection

Fluid samples of cerebrospinal fluid (CSF), blood, and saliva were collected monthly from each of the three experimental goats for seven months. All samples were collected, stored, and analysed following the Alzheimer’s Association Guidelines for handling CSF (Hansson et al., 2021), blood (Verberk et al., 2022), and saliva (Ng et al., 2025). The CSF only was sampled under sedation (0.05 mg/kg xylazine IV). For the lumbar puncture, the skin surface was shaved and scrubbed with alcohol at the lumbosacral joint, and 0.5 ml of 2% lidocaine was injected subcutaneously at the lumbrosacral site. A 20 G 3.5” epidural needle was advanced into the lumbosacral space and approximately 5 ml of CSF was sampled. Blood was collected from the jugular vein into tube containing EDTA anticoagulant and saliva was sampled using cotton swabs inserted into the cheek. All samples were immediately labeled and stored on ice during transport back to the lab within 2 h of collection. To obtain platelet-free plasma, blood was centrifuged for 15 minutes at 2,000 x g. The CSF and plasma samples were then divided into aliquots. After processing, all samples were stored at -80 °C until further use. Before use, saliva samples were thawed at room temperature, and swabs were centrifuged at 13000 RPM at 4 °C for 20 minutes.

### Biomarker analysis

The concentrations of neurodegeneration biomarkers (total tau, phosphorylated tau, GFAP, APP, Aβ_40_, and Aβ_42_) in CSF and plasma were simultaneously measured using the MILLIPLEX^®^ Human neurodegenerative disease panels (Panel 1 for GFAP: HNSS1MAG-95K-01, A-Beta Tau panel: HNABTMAG-68K-04, Panel 4 for S100B: HNDG4MAG-36K-01. Millipore, MA). Only six months of samples were available for GFAP, as some samples were below the kit detection range. NF-M was measured in CSF and plasma by traditional ELISA with a commercial kit (Human Neurofilament Protein M [NF-M] ELISA kit, 76704-648, AFG Bioscience, detection range 3.75-120 ng/ml, sensitivity: 0.39 ng/ml). Assays were performed according to the manufacturer’s instructions, there were no dilutions, and signals were detected at wavelengths specified by the manufacturers using a microplate reader (Cytation 5 Multi-Mode Reader, Agilent, CA).

Presence of Aβ_42_ biomarkers was measured in saliva (n = 21 samples) using an ELISA Sandwich kit (AFG Bioscience Human Ab1-42, Cat. No: 77513-626, detection range: 15.63-1000 pg/ml, sensitivity: 5.31 pg/ml) following manufacturer’s instructions.

### PET/MRI acquisition

A CT scan was performed initially on the goats as a standard precaution to screen for the presence of metallic foreign bodies that could pose a safety risk or contamination to subsequent PET/MRI. For PET/MRI scans, animals were transported from MSU to the University of Alabama at Birmingham’s (UAB) imaging facility, in accordance with standard large animal transport protocols. Animals were housed overnight in the UAB animal facility prior to the procedure to allow for acclimatization and to minimize stress, and were housed after the procedure for post-imaging monitoring and radioactive half-life decay prior to return to MSU. Prior to the procedure, the goats were intubated with a cuffed endotracheal tube and maintained under anesthesia (inhalation isoflurane 2-3% on 100% oxygen) for the duration of the transport and imaging under the guidance of a board-certified anesthesiologist (KF). Goats were sedated with 0.05 mg/kg xylazine and 0.05 mg/kg butorphanol IV for pre-medication and induced with 2-5 mg/kg propofol IV. PET/MRI scans were performed at the start and end of the experiment for longitudinal measurement of neurophysiological and structural changes (Fig. 1). Each goat was positioned prone and administered with a total mean activity of approximately 5 mCi of either [^18^F]DPA-714 (first scan) or [^18^F]-FDG (second scan) via IV catheter and monitored for any adverse reactions. Simultaneous whole-body PET/MRI was acquired dynamically for 60 min (GE sigma PET/MRI) during and post administration of the radiometer. During the scans, optimization of MRI coil selection and imaging sequences were completed to reduce artefacts from the goat’s horns. T_1_- and T_2_-weighted anatomical imaging was acquired with a 64-channel head coil. The initial PET scan used [^18^F]DPA-714 as a marker for TSPO, an indicator of neuroinflammation. Despite previous success in humans (Fang et al., 2022), this marker did not pass the blood-brain barrier in goats due to high plasma protein binding. As the ability for a comparative scan was eliminated, the final PET scan was therefore changed and used [^18^F]-FDG as a marker for glucose metabolism activity. Simultaneous MRI allowed anatomical brain assessment to correspond with PET uptake and activity. MRI scans were reviewed by a radiologist for internal signs of TBI pathology such as acute trauma, regional shrinkage, or microhemorrhage. Qualitative assessment of regional uptake of each radiotracer in the brain from PET imaging was completed.

### Video monitoring

For the length of the experiment, continuous video was captured of the goats from an overhead angle using a ZWO ASI178MM camera set at 15 frames per second and a capture area of 3096 x 2080 pixels, on a black-and-white capture setting. The videos were transferred onto a physical hard drive once a week. Each video was approximately 55 minutes long. Videos that occurred in the night and had no visibility were removed. In total 1206 videos were captured in the dataset, representing daylight hours from October 22^nd^, 2023, to April 12^th^, 2024. 162 files were excluded from this study due to animals being absent for procedures, corrupt files, or dark videos, leaving 1044 total usable videos (Fig. 1). Videos were compressed using VLC (3.0.20 “Vetinari”), sped up 2.5x. and then processed in Adobe Lightroom (v.8.0 x 64) to improve visibility.

### Behavior analysis

Individual headbutts were manually counted from the video dataset. Headbutts were defined as an animal forcefully hitting a conspecific or object with its head. Headbutts were qualified as “high force” if the goats reared up and adopted a bipedal posture before headbutting. For each interaction, the headbutt initiator and receiver was recorded, and both were counted as having received a head impact, except when the receiver was a conspecific’s flank or object, in which case, only the initiator was recorded as receiving a head impact. At the time of submission, manual behavioral counting attained 55% coverage of the total dataset.

### Accelerometer analysis

To quantify the kinematics of headbutting, a 9-axis accelerometer and inclinometer (Witmotion WT901BLECL) was affixed to the back of the horn of goat A. Continuous kinematic data were recorded over twelve discrete tracking sessions, ranging between 30 and 60 minutes in duration. Linear acceleration, angular velocity, and head orientation were measured along three orthogonal axes (x, y, z). Resultant acceleration was calculated as the vector magnitude of the three acceleration axes, providing a measure of overall head motion independent of direction. Consistent with recommendations from the Consensus Head Acceleration Measurement Practices (CHAMP) group, we report both linear and rotational kinematic measures at the head as primary descriptors of impact exposure (Arbogast et al., 2022). Headbutting events were identified from threshold exceedances in resultant acceleration and grouped into discrete events using a temporal separation criterion. To estimate impact forces, the effective mass of the head-neck complex was assumed to be 8% of the animal’s total body mass (∼29.94 kg for goat A), following conservative estimates for ungulates (Loscher et al., 2016). Resultant force was then calculated from resultant acceleration and effective mass and used to characterize the magnitude of headbutting impacts. Resultant angular velocity was calculated from the three gyroscopic axes to quantify rotational head movement during impacts, while maximum head orientation was also recorded from inclinometer measurements to characterize head posture during headbutting events. Additional details of data processing and biomechanical calculations are provided in the supplementary methods.

### Maze analysis

As a measure of cognitive decline, goats were trained to complete a Y-maze, as goats have been shown to be able to perform the task without significant lateralization and retain the skill in the long-term (Langbein, 2012). Each goat performed the Y-maze test weekly over the course of the experiment. If the duration to maze completion was above 5 minutes, the maze run was considered “failed” and excluded from the dataset. Changes in maze completion time was evaluated with a Spearman’s rho correlation test for each goat over time.

### Data analysis & statistics

All data analyses were conducted using R Studio software (Posit team, 2025) using tidyverse() packages. Code and raw data are available on GitHub (https://github.com/NLAckermans/GoatTBI2026).

## RESULTS

### Immunohistochemistry & Immunofluorescence

#### A. Homology of key NDD proteins between human and goat

BLAST analysis confirmed selected biomarkers markers as highly conserved between goat and human (Aβ > 95%, GFAP 95%, S100B 99%, Tau > 90%) (Table 2).

#### B. Neuropils in goat brain

Neuropils were identified in fixed brain tissue of all three adult goats using tau antibodies. Specifically, we detected structures expressing pTau Ser202 as detected by an antibody raised against CP13. We observed neuropil threads expressing pTau interspersed among neuron cell bodies and axons within the prefrontal cortex, and some appeared fragmented with tortuous processes (Fig. 3). These structures were present in similar densities between goats, at 400-900 neuropil threads per goat across a similar volume of tissue. When standardized by tissue volume, this equated to a total of 0.0002 neuropils/mm^3^ for goat A, 0.00002 neuropils/mm^3^ for goat B, and 0.0001 neuropils/mm^3^ for goat C.

Densley clustered neuropil threads expressing pTau were also observed in Goat A around the surface layers (Fig. 2). Clustering of structures expressing pTau at the bottom of a sulcus was measured in the prefrontal cortex of Goat B where all structures expressing pTau were located around a sulcus, and 5/7 were clustered at the bottom of that sulcus (Fig. 2).

**Figure 2.**
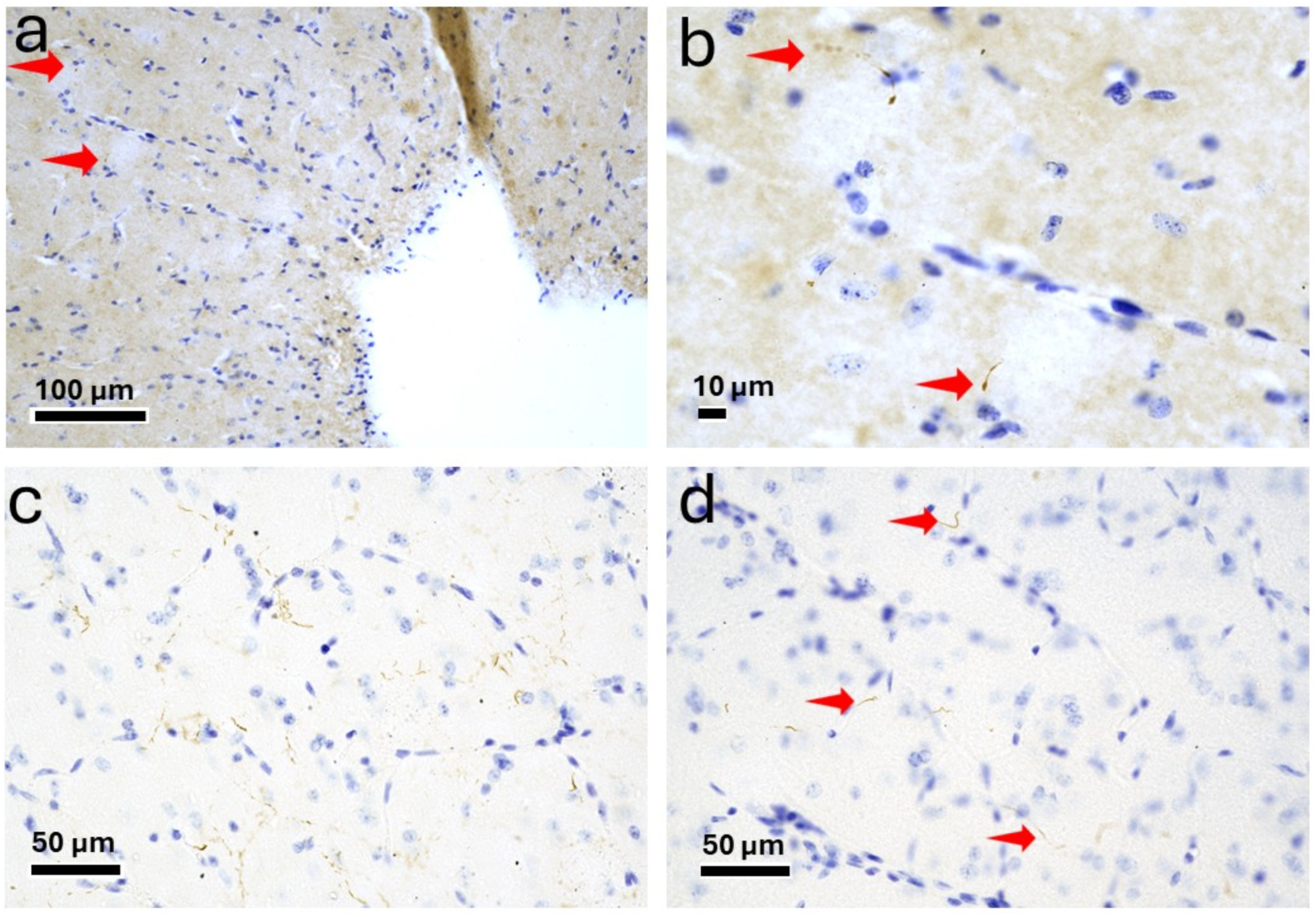
Neuropils expressing abnormally phosphorylated tau in the prefrontal cortex of headbutting goats (*Capra hircus*). Brain tissue of an approx. 1 yr goats. Red arrows non-exhaustively indicate neuropils. pTau in solid dark brown, Nissl-stained cell nuclei in blue. a) Goat B at 20x and b) 40x; c) Goat A, multiple neuropil threads visible in this region; d) Goat C.

We quantified association of structures expressing pTau with blood vessels as in (Ackermans et al., 2022) for goats A, B, and C. All blood vessels > 30 μm in diameter were quantified in one tissue section using Stereoinvestigator’s Optical Fractionator probe, then the percentage of vessels less than 100 μm from a structure expressing pTau, as well as the average distance from the structure were measured and reported. Goat A had 0/23, Goat B had 2/32, and Goat C had 0/14 pTau associated blood vessels.

No structures expressing pTau were detected in either of the younger goats used as comparison, no neurofibrillary tangles were observed in any goats, and no pTau S396/404 expression was detected using an antibody raised against PHF-1.

#### A. Reactive astrocytes in goat brain

Antibodies raised against S100B and GFAP revealed the presence of reactive astrocytes in all three goats (Fig. 3). Furthermore, in all three goats we also detected varicose projection (VP) astrocytes. VP astrocytes are a recently described astrocytic morphology where astrocytes have long, beaded processes, suggested to be related to neuroinflammation and neuropathology (Falcone et al., 2022; Ciani et al., 2025; Ciani et al., 2026). As the VP astrocytes were small, relative to those observed in other species, their presence likely indicates mild pathology in our goats.

**Figure 3:**
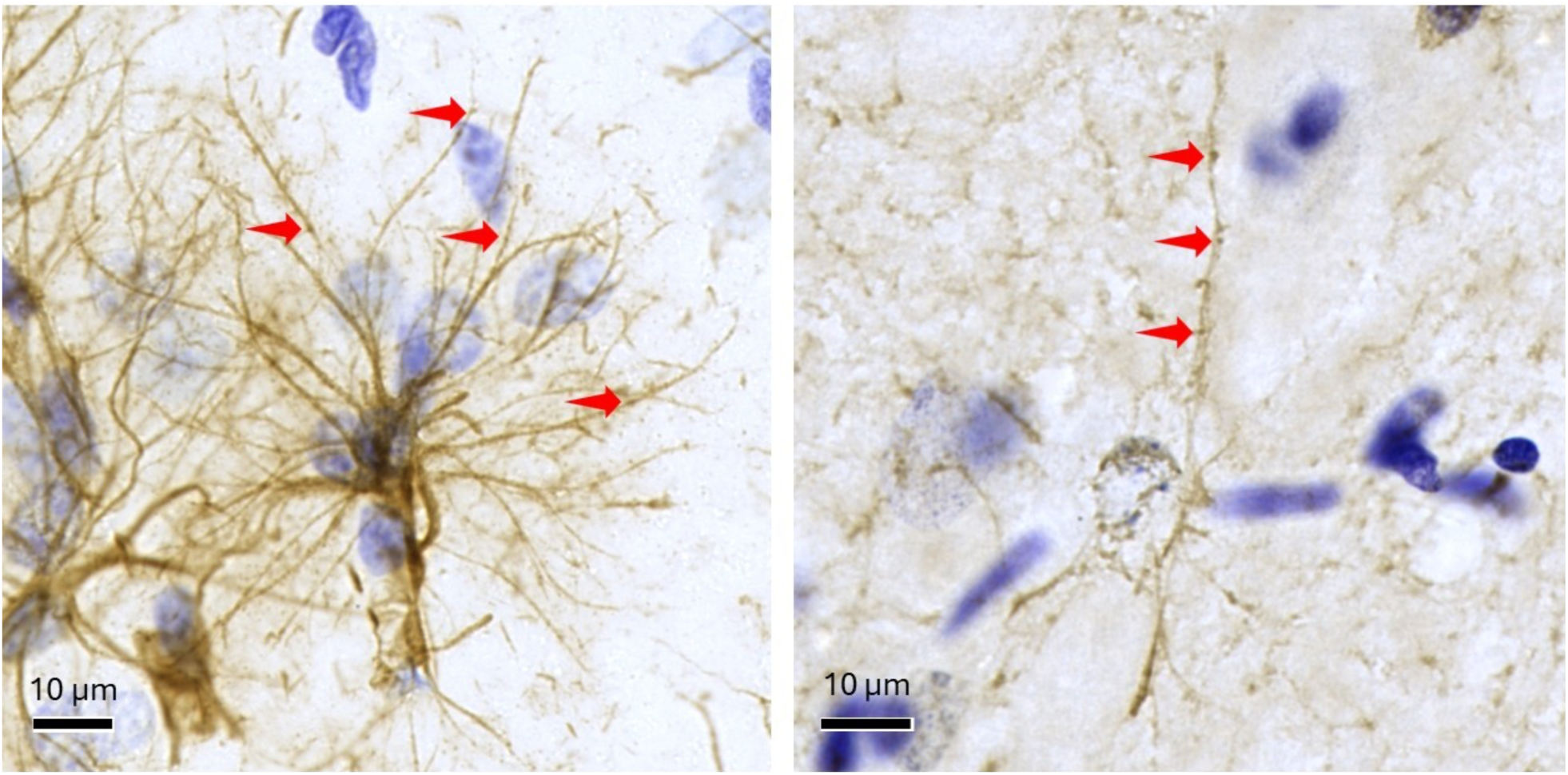
Varicose projection astrocytes in prefrontal cortex of headbutting goats (*Capra hircus*). Brain tissue of approx. 1 yr goats. Varicosities are non-exhaustively marked with red arrows. Left, goat A, astrocytes expressing GFAP. Right, goat B, astrocytes detected with an antibody raised against S100B. Astrocytes in solid dark brown, Nissl-stained cell nuclei in blue.

### Biomarkers

#### CSF biomarkers

The standard curve for the biomarkers ranged as follows: Aβ_42_ (1-1000 pg/ml), Aβ_40_ (10-10000 pg/ml), pTau (1-1000 pg/ml), tTau (10-10000 pg/ml), GFAP (10-100000 pg/ml), S100B (10-10000 ng/ml) and NF-M (3.75-120 ng/ml) (Fig. S1).

#### a. Aβ levels in goat CSF

Aβ_42_ in goat CSF ranged from 382.5 – 1602.3 pg/ml and the average was 881.2 pg/ml. Aβ_40_ in goat CSF ranged from 91.3 – 4447.4 pg/ml and the average was 2345.3 pg/ml. Aβ_42:40_ ratio in goat CSF ranged from 0.2 – 10.2, and the average was 0.8 (Fig. 4).

**Figure 4:**
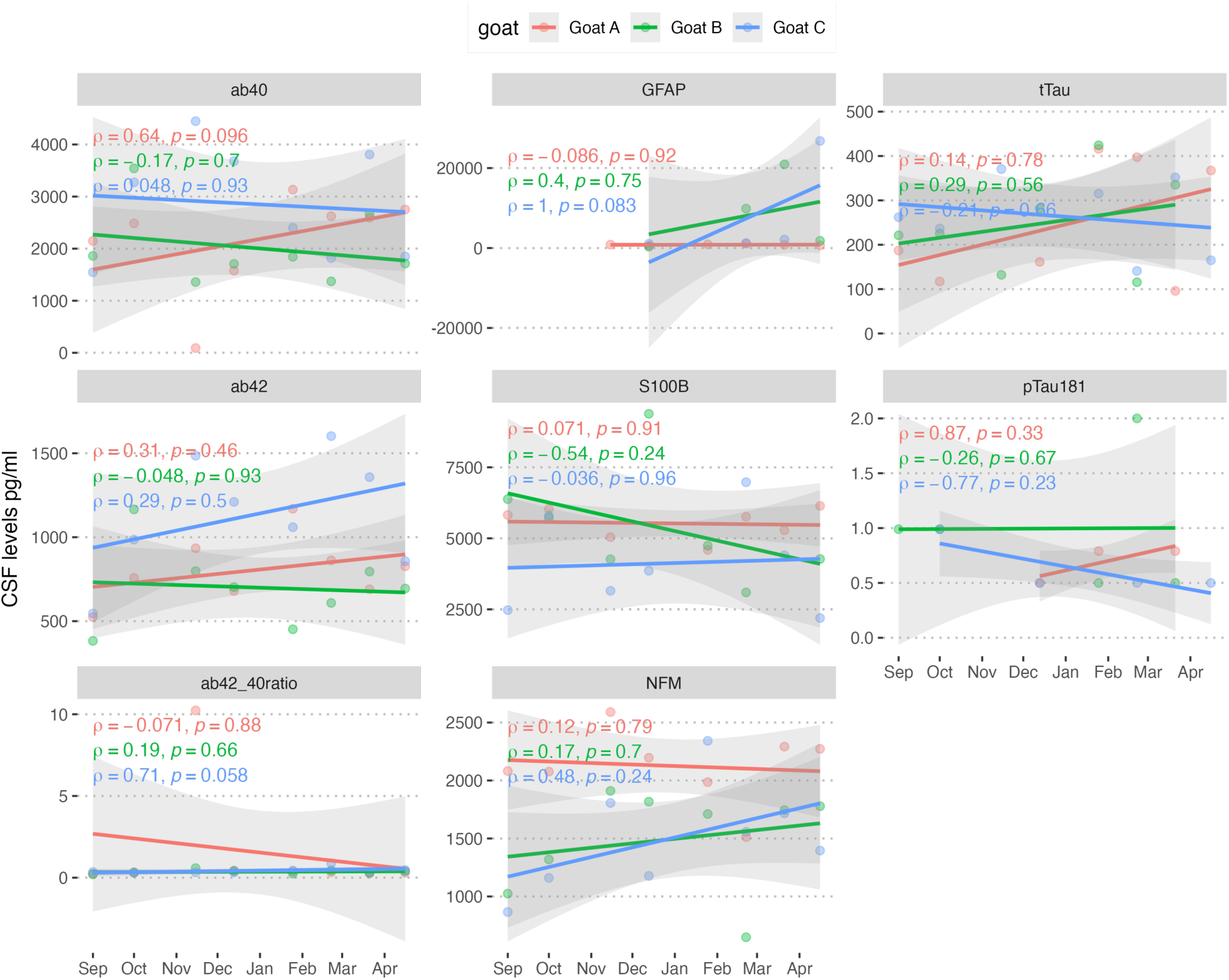
Monthly cerebrospinal fluid (CSF) concentrations of biomarkers in headbutting goats (*Capra hircus*, n = 3). Biomarkers were measured from CSF samples using multiplex ELISA assays and regression lines show the longitudinal trends of the measured biomarkers using Spearman’s rho.

**Figure 5.**
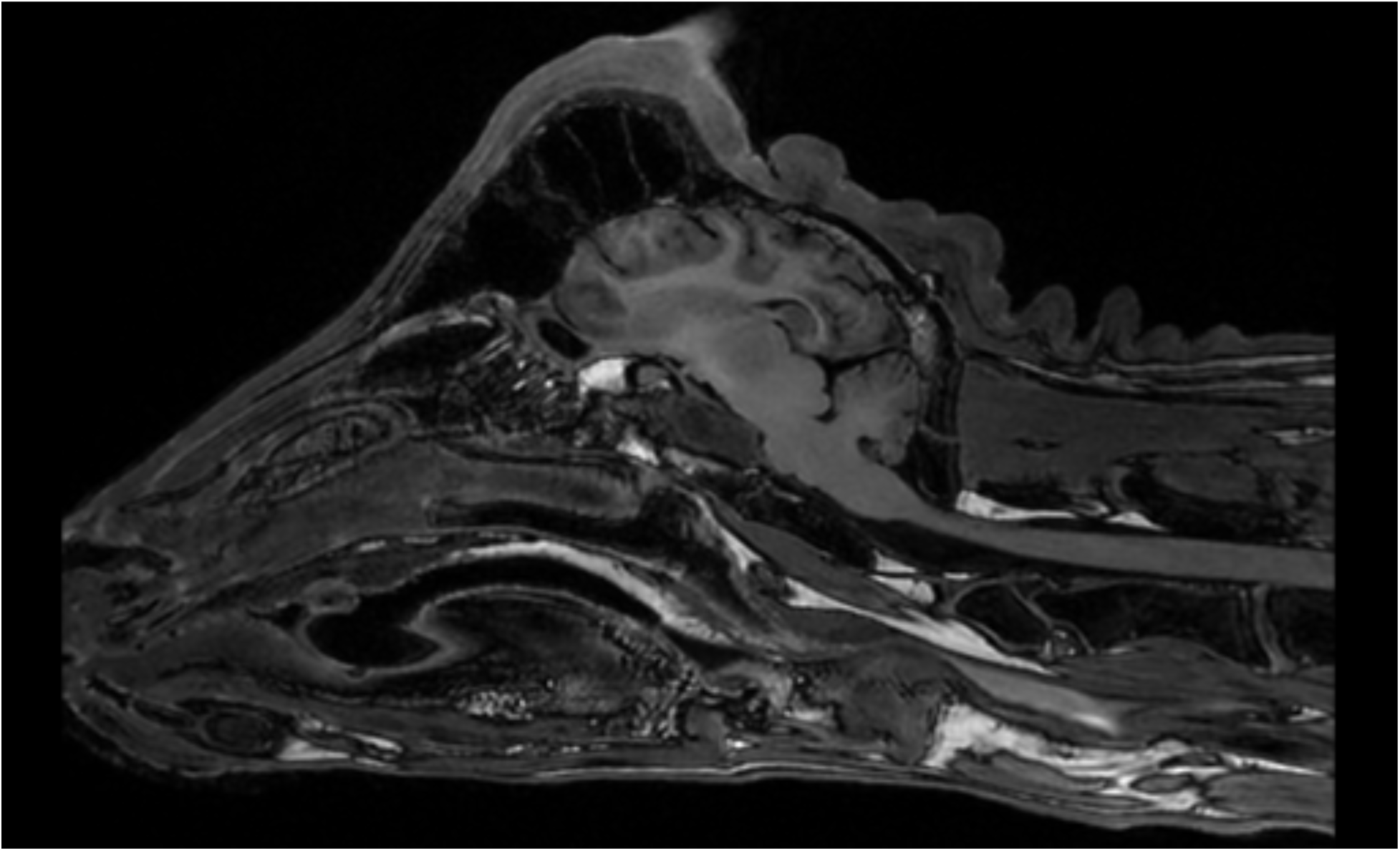
Cross-sectional MRI of a goat (*Capra hircus*) brain. The scan demonstrated no visual changes in anatomical or structural components.

A meta-analysis measured mean CSF Aβ_42_ in AD patients at 183 pg/mL compared with 491 pg/mL in healthy controls using ELISA (Sunderland et al., 2003). Another study considered CSF Aβ_42_ as pathological below 640 pg/ml in humans (Hansson et al., 2007).

In humans, the healthy control Aβ_42:40_ ratio average is 0.21, versus an average of 0.12 in AD patients using ELISA (Spies et al., 2010).

#### b. Total and phosphorylated tau and in goat CSF

Goat CSF tTau ranged from 96 – 424.2 pg/ml, and the average was 253.3 pg/ml. Goat CSF pTau181 ranged from 0.5 - 2 pg/ml, and the average was 0.8 pg/ml (Fig. 4). A meta-analysis found mean CSF tTau in AD patients at 587 pg/mL and at 224 pg/mL in healthy controls using ELISA (Sunderland et al., 2003). The mean CSF pTau181 was 95 pg/ml in AD patients and 48 pg/ml in controls (Spiegel et al., 2015).

#### c. GFAP in goat CSF

GFAP in goat CAF ranged from 344.9 – 26,780.6 pg/ml, and the average was 4,932.7 pg/ml. In the last three months of the experiment, multiple measures for Goats B and C were above the threshold for healthy GFAP CSF levels in humans, ranging from 1,800 to 26,800 pg/ml (Fig. 4). The range for human GFAP range in healthy controls is 100-1,300 pg/ml (Rosengren et al., 1994), whereas the average for human AD is 3,200 pg/ml (Ishiki et al., 2016).

#### d. S100B in goat CSF

S100B in goat CSF ranged from 2,196 - 9,383 pg/ml, and the average was 5,019 pg/ml (Fig. 4). The range for human S100B healthy controls is 1600-3300 pg/ml (Hayakata et al., 2004; Hajduková et al., 2015), whereas pathological human S100B range (TBI) is 63,0000 pg/ml on average (Hayakata et al., 2004).

#### e. NFM in goat CSF

NFM in goat CSF ranged from 648.49 – 2,590.25 pg/ml, and the average was 1,707.65 pg/ml (Fig. 4). The range for human NFM in healthy controls is 230 pg/ml on average, whereas pathological human range (stroke) is 6,000-11,000 pg/ml (Martínez-Morillo et al., 2015a).

**Table 7.** CSF biomarker levels in freely headbutting goats (*Capra hircus*) over eight months. Biomarker levels in pg/ml, values outside the detection range were excluded.

| Goat | Date | ab42 | ab40 | ab42:40 | tTau | pTau181 | NFM | S100B | GFAP |
| --- | --- | --- | --- | --- | --- | --- | --- | --- | --- |
| Goat A | 2023-09-01 | 524.28 | 2146.56 | 0.24 | 187.02 |  | 2081.82 | 5822.9 |  |
|  | 2023-10-01 | 757.49 | 2485.45 | 0.30 | 117.26 |  | 2076.56 | 6033.13 |  |
|  | 2023-11-15 | 934.31 | 91.31 | 10.23 |  |  | 2590.25 | 5041.74 | 841.80 |
|  | 2023-12-13 | 679.88 | 1576.22 | 0.43 | 161.39 | 0.50 | 2196.04 |  | 553.97 |
|  | 2024-01-25 | 1169.37 | 3133.01 | 0.37 | 416.05 | 0.79 | 1986.20 | 4591.08 | 917.31 |
|  | 2024-02-22 | 861.86 | 2623.21 | 0.33 | 397.32 |  | 1511.49 | 5759.33 | 1108.19 |
|  | 2024-03-21 | 690.15 | 2591.94 | 0.27 | 96.02 | 0.79 | 2292.27 | 5282.64 | 833.12 |
|  | 2024-04-16 | 827.22 | 2751.49 | 0.30 | 367.33 |  | 2272.20 | 6142.44 | 669.65 |
| Goat B | 2023-09-01 | 382.52 | 1860.52 | 0.21 | 221.22 | 0.99 | 1024.60 | 6378.35 |  |
|  | 2023-10-01 | 1165.76 | 3541.45 | 0.33 | 225.31 | 0.99 | 1319.27 | 5723.83 |  |
|  | 2023-11-15 | 798.07 | 1359.9 | 0.59 | 132.18 |  | 1910.54 | 4263.72 |  |
|  | 2023-12-13 | 703.01 | 1708.13 | 0.41 | 281.6 |  | 1817.22 | 9382.58 | 344.85 |
|  | 2024-01-25 | 451.71 | 1843.81 | 0.24 | 424.21 | 0.50 | 1711.16 | 4740.01 |  |
|  | 2024-02-22 | 608.55 | 1372.72 | 0.44 | 115.52 | 2.00 | 648.49 | 3096.29 | 9852.35 |
|  | 2024-03-21 | 795.17 | 2666.96 | 0.30 | 335.24 | 0.50 | 1744.84 |  | 20943.92 |
|  | 2024-04-16 | 694.51 | 1713.07 | 0.41 |  |  | 1778.35 | 4273.39 | 1802.85 |
| Goat C | 2023-09-01 | 545.8 | 1545.99 | 0.35 | 262.04 |  | 866.26 | 2473.21 |  |
|  | 2023-10-01 | 985.71 | 3273.38 | 0.30 | 236.54 | 0.99 | 1160.29 | 5799.68 |  |
|  | 2023-11-15 | 1485.72 | 4447.41 | 0.33 | 370.6 |  | 1806.14 | 3153.5 |  |
|  | 2023-12-13 | 1210.31 | 3680.18 | 0.33 |  | 0.50 | 1177.99 | 3856.98 | 1060.94 |
|  | 2024-01-25 | 1059.77 | 2400.55 | 0.44 | 315.25 |  | 2342.00 |  |  |
|  | 2024-02-22 | 1602.32 | 1817.05 | 0.88 | 140.92 | 0.50 | 1557.57 | 6975.7 | 1241.92 |
|  | 2024-03-21 | 1358.15 | 3807.02 | 0.36 | 352.01 |  | 1716.78 | 4402.85 | 2106.06 |
|  | 2024-04-16 | 856.93 | 1850.28 | 0.46 | 164.98 | 0.50 | 1395.35 | 2195.57 | 26780.57 |

Ab42 = amyloid-beta 42, ab40 = amyloid-beta 40, tTau = total tau, pTau181 = phosphorylated tau 181, NFM = neurofilament light, GFAP = glial fibrillary acidic protein

### Plasma biomarkers

#### a. Aβ levels in goat plasma

Aβ_42_ in goat plasma ranged from 4.7 – 23.7 pg/ml, and the average was 12.9 pg/ml. Aβ_40_ in goat plasma ranged from 69.6 – 151.7 pg/ml, and the average was 122.36 pg/ml. Aβ_42:40_ ratio in goat plasma ranged from 0.04 – 0.17, and the average was 0.11 (Fig. S1).

Mean AD human plasma Aβ_42_ was 68.7 pg/ml, as compared to healthy controls at 58.8 pg/ml, mean AD human plasma Aβ_40_ was 153.6 pg/ml, as compared to healthy controls at 133.3 pg/ml, and mean AD human plasma Aβ_42:40_ ratio was 0.49, as compared to healthy controls at 0.48 (Mayeux et al., 2003).

#### b. Tau and phospho-tau181 in goat plasma

There was no signal for tTau in goat plasma. However, pTau181 in goat plasma ranged from 0.5 – 1.27 pg/ml, and the average was 0.93 pg/ml. Mean AD human plasma pTau181 was 0.17 pg/ml as compared to a healthy control cohort at 0.04 pg/ml (Tatebe et al., 2017).

#### c. GFAP in goat plasma

There was no signal for GFAP in goat plasma.

#### d. S100B in goat plasma

S100B in goat plasma ranged from 40.2-119.9 pg/ml, and the average was 75.6 pg/ml. In the literature, the level for healthy control goat S100B plasma was 440 pg/ml on average, whereas the pathological range (*C. cerebralis* infection) was 870 pg/ml on average (Uztimür and Dörtbudak, 2023). In addition, plasma S100B was generally elevated in TBI patients as compared to healthy controls (exact values not specified) (Chen et al., 2019).

#### e. NFM in goat plasma

NFM in goat plasma ranged from 877.92 – 1699.89 pg/ml, and the average was 1095.8 pg/ml. While plasma ranges for human NFM are infrequent in the literature, the NFM range/levels for healthy control humans is 0.26-8.57 pg.ml with a median of 2.29 pg/ml, as compared to a range of 3.48-45.4 with a significantly higher median of 13.3 for pathological human (TBI) (Martínez-Morillo et al., 2015b).

**Table 8.**
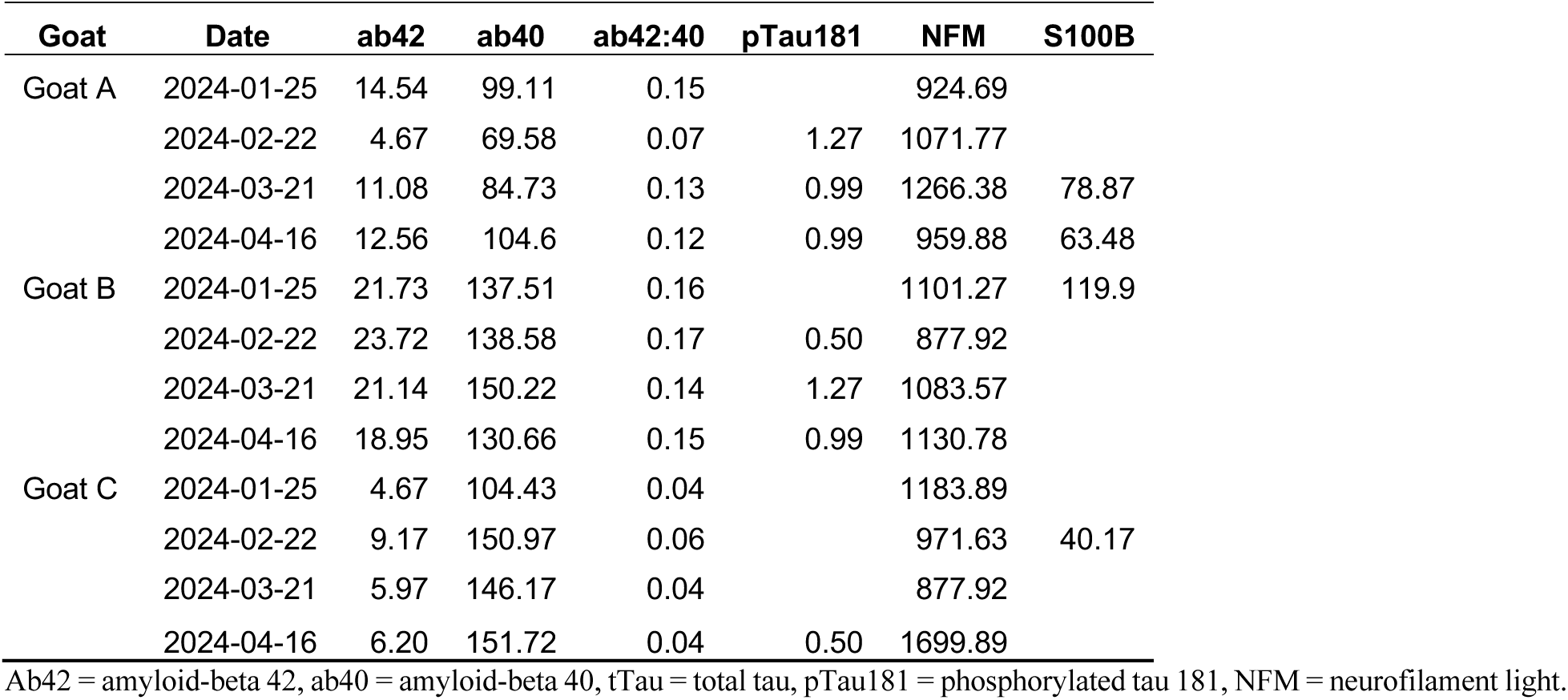
Plasma biomarker values in headbutting goats (*Capra hircus*) over four months. Biomarker values in pg/ml, samples outside of the detection range were excluded.

### Aβ in goat Saliva

Levels of Aβ_42_ in goat saliva ranged from 22.4 – 47.9 pg/ml, average 33.6 pg/ml. The mean Aβ_42_ level for healthy control human saliva is 21.1 pg/ml, and the mean for pathological human saliva (AD) is Aβ_42_ 51.7 pg/ml (Sabbagh et al., 2018). In our study, of the 21 samples tested, only 6 had adjusted optical density values within detection range.

**Table 9.** Ab_42_ biomarker measured in saliva of freely headbutting goats over seven months (*Capra hircus*, n = 3). Biomarker values are in pg/ml, metrics outside the detection range were not reported.

| Goat | Date | ab42 |
| --- | --- | --- |
| Goat A | 2024-01-25 | 34.59 |
|  | 2024-02-22 | 22.35 |
|  | 2024-04-16 | 22.35 |
| Goat B | 2024-02-22 | 42.67 |
| Goat C | 2023-11-15 | 31.68 |
|  | 2024-01-25 | 47.86 |

Ab42 = amyloid-beta 42

#### PET/MRI

None of the specimens showed any external brain trauma and MRI scans were reviewed by a radiologist for internal signs of TBI pathology such as acute trauma, regional shrinkage, or microhemorrhage. No such alterations were observed in our specimens (Fig. 3).

At the initial imaging timepoint (goats at 6 mo old), [^18^F]DPA-714, a radiotracer for TSPO imaging, did not pass the blood brain barrier in either goat, eliminating the prospect for a comparative PET scan at the second imaging timepoint. [^18^F]FDG was used at timepoint two as a mixed marker for glucose metabolism and inflammation was successfully imaged in the goat brain for the first time. For FDG-PET/MRI, regional differences appeared between the FDG uptake on the three goats on PET, but with no corresponding anatomical variations assessed via MRI. All goats showed heightened frontal-dominant FDG uptake, and one displayed an additional asymmetrical uptake, which is often detected in clinical early stages of AD (Mosconi, 2005). Further experiments are required to investigate the underlying biological mechanism of this increased uptake as FDG is a nonspecific marker of glucose transport.

### Behavior

#### Headbutting

Video analysis revealed that each goat sustained 227 head impacts daily on average, with each goat sustaining approximately between 5,000 and 8,000 impacts over the course of the experiment, of which 8% were considered high force headbutts (Fig. 1). Goat B accumulated the most headbutts of all three goats (7745 impacts), and 22% of those impacts were against the fence, as opposed to another goat. Goat C headbutted the second-most (5596 impacts, 99% on other goats), and Goat A headbutted the least (5170 impacts, 97% on other goats) (Table 10).

**Table 10.**
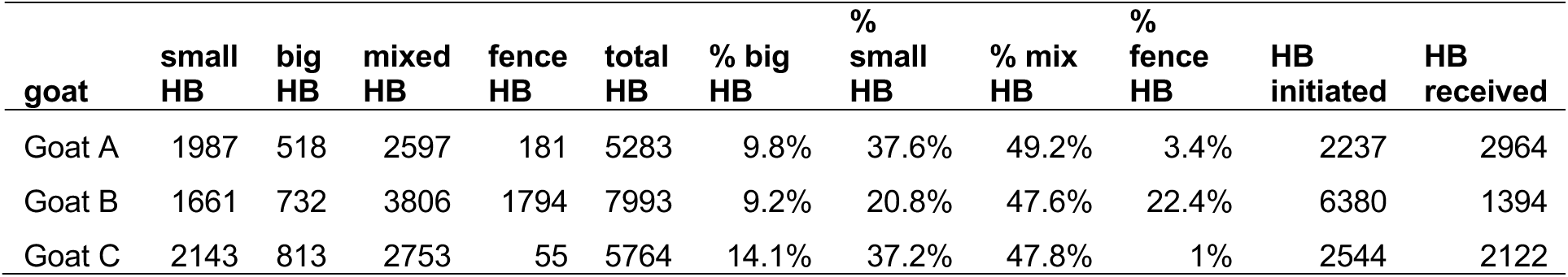
Cumulative headbutt (HB) metrics for each goat over a six-month pilot experiment.

Goat B initiated headbutts 2-4 times more frequently than goats A and C (4,000 times compared to 1,000 and 2,000, respectively). Similarly, goat A received about half as many headbutts than the other two (1,000 and 2,000 respectively).

### Accelerometers

Accelerometer recordings identified between 12 and 372 discrete high-impact events per tracking session, with an average impact duration of 0.03 to 0.18 s per event (Table S1). Baseline, non-impact activity produced low average kinetic profiles, with mean resultant acceleration ranging from 0.91 to 1.31 g (8.92 to 12.85 m/s^2^) and mean estimated forces ranging between 21.43 and 30.8 N. In contrast, headbutting behaviors generated pronounced kinetic profiles, with maximum resultant accelerations reaching 16.53 g (162.08 m/s^2^), and estimated impact forces reaching 388.18 N (Fig. 6, S3-S4). Peak resultant angular velocities reached 3463.5° s⁻¹ (60.45 rad/s). Cumulative impact impulse was highest on the most active day, reaching 98,676.30 N·s, indicating repeated loading of the head-neck complex across extended observation periods.

**Figure 6.**
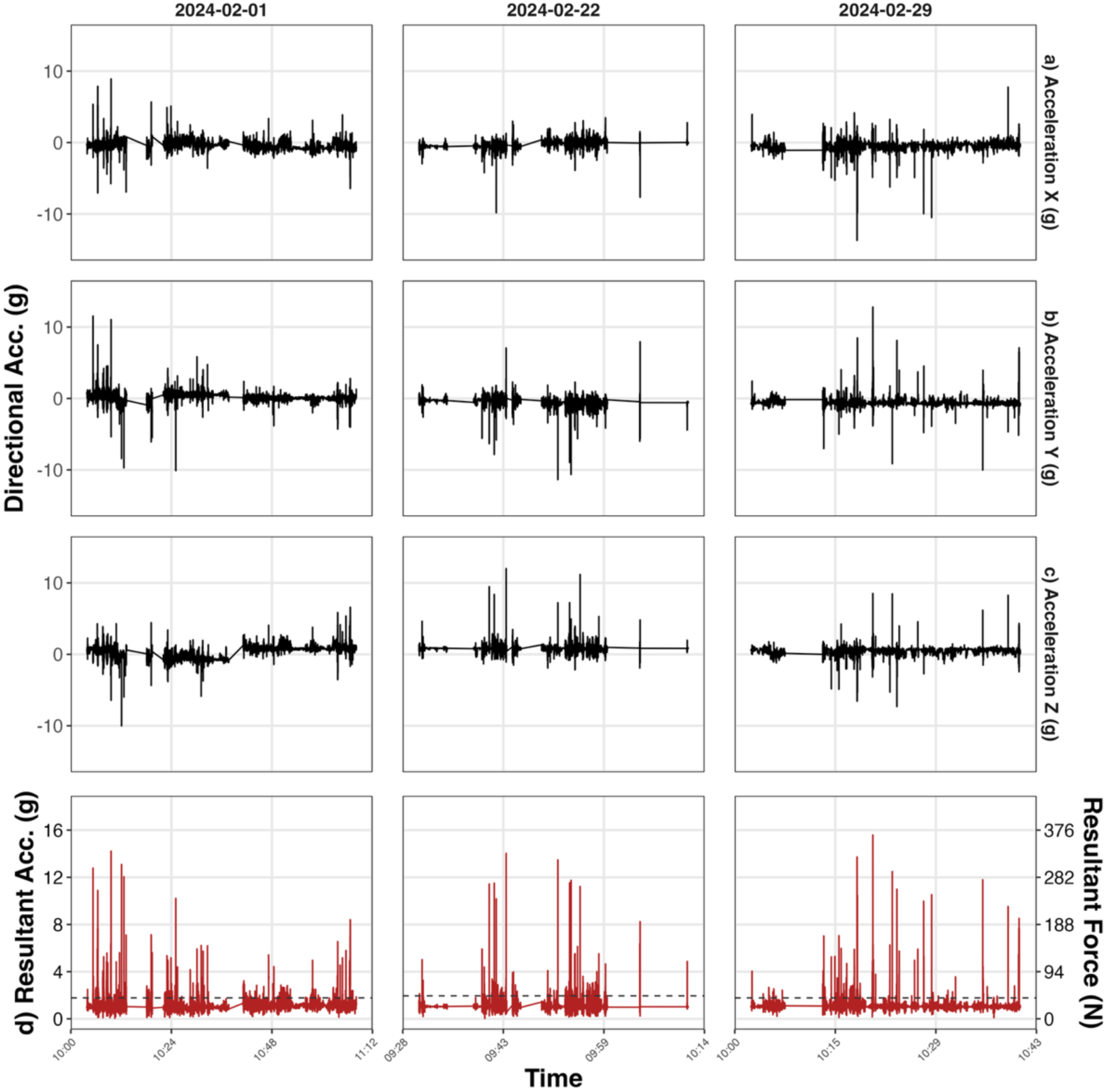
Triaxial acceleration and resultant loading during naturally occurring headbutting events in a domestic goat (*Capra hircus*). Directional acceleration measurements along the x-, y-, and z-axes are shown for the three selected recording sessions with the greatest number of detected high-impact events for goat A (top panels; a-c). Corresponding resultant acceleration magnitude (in g-force) and estimated resultant force (in Newtons) are shown below for each session (d).

### Cognition test maze

Time to complete the Y-maze, as well as branch choice remained stable over six months for all goats (Fig. 7), indicating neither significant improvement nor loss of cognitive function as per a Spearman’s Rho test. Retaining stable memory of the maze over multiple months is consistent with the literature (Langbein, 2012).

**Figure 7.**
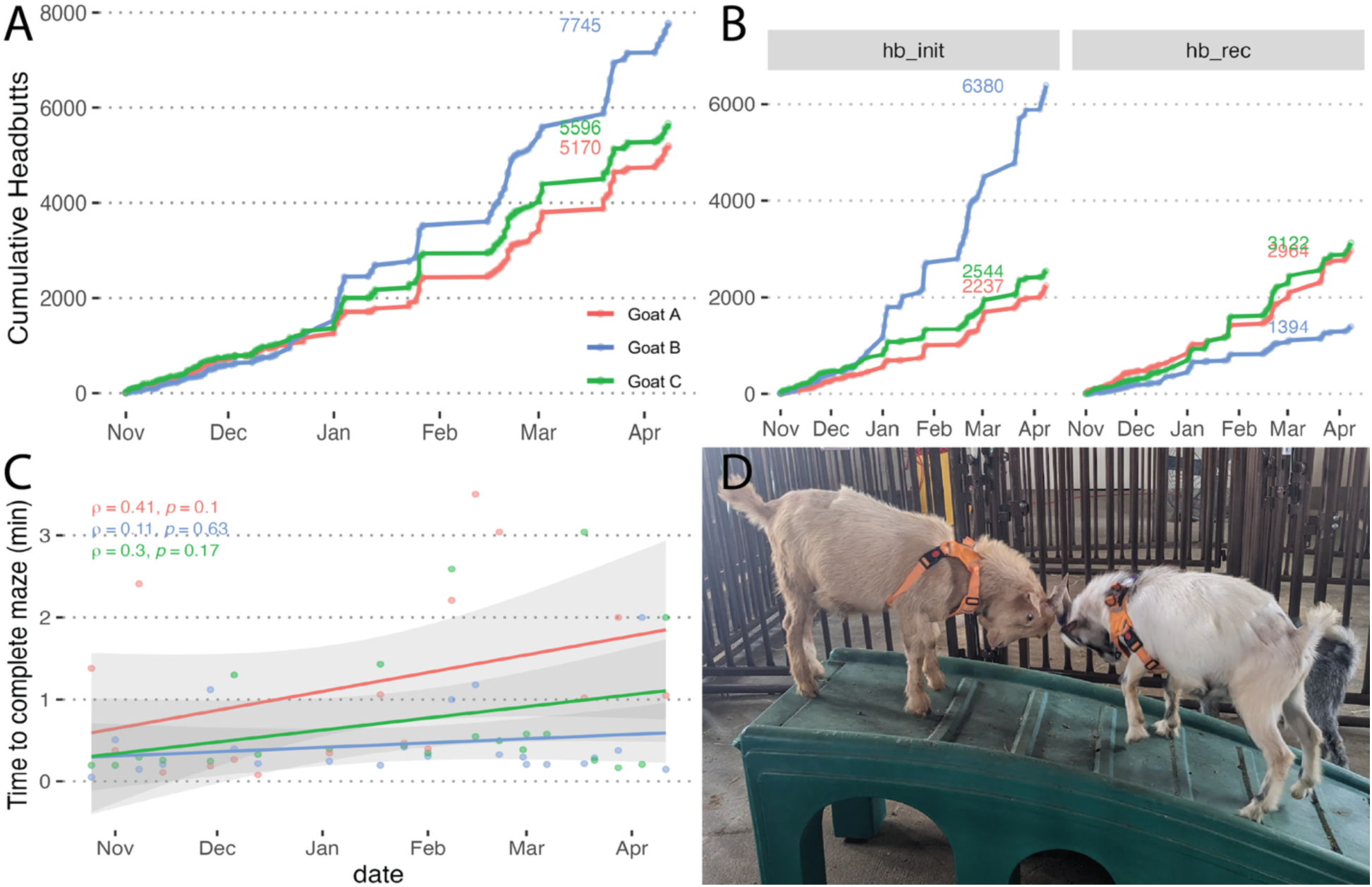
Metrics of three headbutting goats (*Capra hircus*) over six months. A: Number of headbutts in three freely headbutting goats over seven months, 68% coverage. B: Cumulative headbutts per goat, faceted by initiator or receiver. C: Maze completion time for individual goats across the experimental period. D: Goat A and goat C headbutting.

## DISCUSSION

We observed freely headbutting goats over six months and detected multiple markers of neurodegeneration during the experiment as well as postmortem. Hyperphosphorylated neuronal tau was observed in the prefrontal cortex of all goats, alongside reactive-state VP astrocytes in pathological conformations. In complement, CSF levels of Aβ_42_, GFAP, and NF-M in our goats were outside of the healthy range for humans, indicating increased amyloid protein buildup, gliosis, and axonal damage. While behavior remained stable across the experiment, each goat sustained around 200 head impacts per day at approximately 10 g, accumulating over 5,000 headbutts each, making this study the first to directly quantify headbutting rates in any animal. In sum, our results support the novel notion that goats self-inflict brain injury from headbutting, as measured by biomarker and histopathology analysis. Goats and other headbutting domestic bovids likely face a similar tradeoff to their wild counterparts, balancing reproductive success against chronic brain damage (Ackermans, 2023). This study encourages the use of goats as a translational model for the study of acute TBI and its longitudinal development into chronic neurodegenerative disease.

### Immunohistochemistry

Phosphorylated tau aggregates were observed in the prefrontal cortex of all three goats in the form of neuropil threads. The number of neuropil threads expressing pTau varied only slightly between goats and was overall relatively low, and none were observed in the younger comparison goats < 1 yr. This study is the first to observe such structures in goats, although Davies et al. (2022) had previously identified fibers and plaques expressing pTau in 1-2 yr sheep. In Artiodactyla, tau aggregation begins in neurites and axons before spreading to neuronal somas (Nakayama et al., 2025). In our samples, no structures expressing PHF-1 were detected and previous studies in bovids had only detected PHF-1 in animals older than 5 yr (Table 2). In humans, antibodies raised against PHF-1 detects a 4R/3R Tau isoform at intermediate NFT maturity levels, whereas antibodies raised against CP13 detect early maturity 4R tau pretangles (Moloney et al., 2021). Thus, successful CP13 detection as opposed to PHF-1 detection in the present study indicates presence of more 4R than 3R tau and taken with age and tau aggregation stage, our results indicate an early stage of neurodegeneration in these goats. Considering the young age of our goats, ∼1 yr which would approximate an 18 year old human or a two year old rat (Carr et al., 2026), it is unlikely that this result is attributable to age-related neurodegeneration. In addition, the lack of pTau in the 3- and 8-month-old goat comparisons indicates that pTau likely begins accumulating after 8 months in these animals. In the literature, structures expressing pTau are much more numerous in artiodactyls over 5 years of age and severity increases with age (Nakayama et al., 2025), to the extent that pTau neuropil threads were described as “very numerous” in aged sheep (Reid et al., 2017), and were orders of magnitude in older muskoxen, when quantified using the same method as the present study (Ackermans et al., 2022) (Table 2). Thus, while pTau appears to accumulate naturally with age in artiodactyls, accumulation severity may be affected by frequency and force of headbutts sustained over an individual’s lifetime. These two factors are difficult to disentangle, as increasing headbutts and age occur simultaneously and precise headbutting numbers require more investigation.

In humans, neuropil threads are reported to constitute 85-90% of cortical tau pathology (Mitchell et al., 2000), and clinically, they are more consistently associated with repetitive, as opposed to single head traumas (Omalu et al., 2005; Armstrong et al., 2017). However, pTau can occur clinically as early as childhood and in young adults (Braak and Del Tredici, 2011) and low rates of neuropil threads expressing tTau have been observed in non-TBI control brains < 20 yr (Smith et al., 2003). Tau phosphorylation is also thought to play a role in promoting axonal growth in the developing brain, however, it is unclear whether the developmental phosphorylation pattern is different from that of pathological pTau plaques (Hefti et al., 2019). Species differences in development and tau phosphorylation add another layer of complication to the tauopathy puzzle. On their own, a low number of neuropil threads expressing pTau in our samples do not solely confirm impact-related neuropathology. However, TBI and chronic neuropathology are defined by a combination of factors, and in our specimens the presence of pathological pTau neuropil threads concentrated in the sulcal depths, as opposed to an even dispersion, indicates trauma-induced neuronal damage (McKee et al., 2023) as opposed to solely inflammation- or age-related neuropathology.

In addition to the specific pTau pattern, another indicator of neuropathology in our samples is the presence of reactive-state astrocytes expressing GFAP, a characteristic morphological change displayed in human TBI and AD (Anderson et al., 2013; De Sousa, 2022), as astrocytes produce more GFAP and S100B under stress (Donato et al., 2013; Michetti et al., 2019; Leipp et al., 2024).. Furthermore, VP astrocytes, a recently described pathological astrocyte morphology in non-human species (Falcone et al., 2022; Ciani et al., 2025; Ciani et al., 2026) were apparent in all three goats, an indicator of mild neuroinflammatory pathology. Together, the immunohistochemical markers indicate an early state of neuronal damage in the headbutting goats.

### Fluid biomarkers

The presence of pathological markers in the brain tissue was also reflected in the fluid biomarkers. In the goat’s CSF, GFAP levels were elevated, with late timepoints showing levels were within the range previously measured in humans AD (Chatterjee et al., 2021; Marksteiner and Humpel, 2025).

A decrease in CSF Aβ_42_ is part of the ATN classification for AD (Jack Jr et al., 2018), and some of the CSF Aβ_42_ measures in our goats were below the value considered pathological in humans (Andreasen et al., 1999). However, in the absence of a reference range for goats, our measured Aβ_42_ range could be interpreted as neuropathological or physiological. The Aβ_42:40_ ratio is considered a more reliable indicator of amyloidogenic shifts than the individual absolute levels of Aβ_42_ or Aβ_40_ alone (Amft et al., 2022), and in our goats, this metric was above healthy human range, when values below would be considered pathological. Our results may reflect increased overall Aβ peptide production with little aggregation, or different enzymatic cleavage patterns of APP, or species-specific differences in APP and Aβ metabolism (Xu et al., 2015; Kimura et al., 2016).

The measured CSF tTau in our goats is similar to values published for human AD (543 pg/ml) although the goats’ pTau concentration was quite low relative to the tTau, possibly reflecting lower phosphorylation rates compared to that recorded for human AD (Baldeiras et al., 2018).

While blood and saliva samples provided some results, indicating that human-target ELISA kits can detect biomarkers in goat samples, the incomplete data did not permit longitudinal biomarker detection, therefore we focused on CSF markers.

Overall, while no statistical trends were present, multiple neurodegeneration-specific biomarkers were recorded outside of the healthy range in our headbutting goats. Severe neurodegeneration was not expected in 1 yr goats, however, combining pathological criteria for CSF biomarkers alongside the pathology detected by IHC, our results indicate the goats already demonstrate the criteria for mild brain damage at this age.

#### PET/MRI

PET/MRI revealed that all goats showed more frontal-dominant FDG activity, and one displayed an additional asymmetrical activation, which is often detected in clinical early stages of AD (Mosconi, 2005). While hypometabolism is an indicator of late-stage AD (Chételat et al., 2020), regional increased FDG uptake could be indicative of localized inflammation in the prefrontal cortex in response to continuous head impacts. Quantitative regional standardized uptake values (SUV) of FDG in additional subjects could provide additional characterization and understanding of inflammatory signaling following continuous head impacts.

### Video analysis and behavior

Over the course of the experiment, each goat accumulated 5,000 to 7,000 head impacts as measured by video analysis. Future studies can build on this example to acquire objective measurement of biomechanical exposure to headbutts and assign an injury burden score to each individual, allowing for a dose-response relationship between head impact and neuropathological phenotypes.

Behavior measurements indicated that the goats completed the Y-maze successfully, with stable latency throughout the experimental period. In one instance (1/11/24) discarded from the analysis, the maze was run shortly after recovery from anesthesia and all animals showed much higher latency, reinforcing the maze as a positive measure of impairment in goats. An in-depth behavioral analysis, as well as automated behavior tracking will be the focus of future publications. Severe behavioral changes were not expected in these young animals, and the video analysis proved successful in quantifying individual impacts.

### Accelerometer

Accelerometer recordings of headbutting generated repeated exposure to substantial cranial loading in both linear and rotational components. The recorded peak accelerations and impact forces indicated the transmission of substantial mechanical loads through the skull during intraspecific interactions. Peak resultant accelerations (162.08 m/s^2^ or ∼16.5 g) impact forces (388 N) fall within the range reported for repetitive, non-concussive head impacts in human athletic activities, particularly soccer heading and contact sports (Basinas et al., 2022). This suggests that naturally occurring goat headbutting provides a useful comparative model for investigating the long-term consequences of repetitive head impacts. Similar combinations of repetitive moderate impacts and high-magnitude events have been documented in human contact sports and other head-impact systems and are increasingly recognized as an important contributor to cumulative mechanical loading and long-term injury risk (Post and Blaine, 2015; Daneshvar et al., 2023). In vertebrate skeletal tissues, repetitive loads are known to drive strain-dependent biological responses, including localized bone remodeling and structural reinforcement in frequently loaded regions (Frost, 1994, 2003; Christen et al., 2014). These structural changes suggest that the loading environment quantified in the present study provides a meaningful mechanical context for interpreting histological and morphological analyses of cranial tissues. The peak resultant angular velocities (60.45 rad/s) recorded in the present study indicate that headbutting interactions cannot be interpreted or modeled solely as linear, one-dimensional impact events, but rather that intense rotational loading routinely accompanies linear acceleration. Indeed, rotational kinematics are strongly associated with complex tissue deformation within cranial systems as a primary determinant of neural and deep connective tissue strain in both human and experimental animal models (Post et al., 2017; Lota et al., 2022; Daneshvar et al., 2023). While this study did not directly measure internal strain or tissue deformation in real time, the high-velocity angular waveforms and their tight coupling to peaks in linear acceleration predict that the goat skull is able to withstand multi-axial, torsional loading profiles during headbutting. To withstand catastrophic failure, multiple cranial structures likely contribute to stress distribution during headbutting. In particular, cranial sutures act as strain-dissipating interfaces during high-impacts (Srinivas et al., in prep).

### Limitations

Given the novel nature of this work, this study was subject to various limitations that point to areas of improvement. This pilot study was limited to n = 3 goats of the same age and sex, for which we did not have life history data before their acquisition. Future studies with larger, more diverse cohorts in sex and age are warranted to increase translatability. This being an observational study, there was no negative control group, however, reproducibility is based on objective measurement of biomechanical exposure and phenotypic outcome, as opposed to the assumption that all head impacts are equivalent. Preventing goats from headbutting is logistically complicated, as goats of all ages and sexes, as well as de-horned goats headbutt, and isolation elevates stress levels and can increase headbutting. While providing physical protection of the head has been suggested, the actual effect on the reduction of brain trauma would require its own controlled study.

Regarding IHC results, it is difficult to directly compare rates of pathological occurrences to the literature, owing to a lack of standardization of reporting methods. Examples include a given number of structures per section but omitting section volume; and using simple descriptors such as “very numerous”, or “multiple structures”. We suggest future studies adopt a metric of [estimated population]/mm^3^ or, at the least [number of structures] per mm^3^ of brain tissue to increase comparative prospects. Future studies on this model would benefit from investigating additional cell types in more depth. Microglia changes with aging have recently been identified in sheep (Carr et al., 2026), and pathological astrocytic morphologies like VP astrocytes have yet to be investigated in many species.

Regarding biomarkers, high levels of inflammatory markers in blood and CSF can be caused by infectious CNS disease and parasitic infection (Aminlari and Mehran, 1988; Schöb et al., 2023; Uztimür and Dörtbudak, 2023), however, in our study trauma-related brain damage pathology is distinguished from infectious damage or normal aging by the co-occurrence of markers for inflammation as well as neurodegeneration patterns expressing pTau in brain tissue (McKee et al., 2023). More frequent longitudinal CSF sampling in this study may have provided view of biomarker dynamics, however, this method was not recommended by our veterinary team due to the possibility of increasing scarring in the lumbo-sacral region. In future studies, the use of a more precise protein assay method with a larger dynamic range such as would improve biomarker detection and precision of fluid biomarker measurements (Hansson et al., 2021; Schindler et al., 2024b). It should also be noted that, due to an experimental error, blood samples for the first two months were rendered unusable for biomarker analysis, and low saliva volume led to non-standardized saliva sample volume. These issues resulted in inconclusive plasma and saliva data, but the data remain valuable as a proof of concept using human-targeted techniques in our goat model.

As TSPO-PET has previously been successfully applied to rodents and humans (Guilarte et al., 2022), it was an unexpected finding that it did not pass the blood-brain barrier in the goat model. Additionally, with the lack of biological information extracted from the TSPO PET, the pilot study required switching of the radiotracer for timepoint two, thereby eliminating a comparative longitudinal study. However, FDG PET provided preliminary noninvasive spatial information on inflammatory signaling, demonstrating success of PET/MRI in this model. Further investigation of PET tracers related to neuroinflammation and NDD in this model is warranted.

In the video analysis, 100% coverage of all headbutts was not possible due to time constraints. However, the current coverage provides sufficient insight on headbutting in relation to biomarkers. A more comprehensive overview of individual behavioral dynamics is in progress. In the present study, two accelerometers were lost, presumably ingested by the goats, and acquiring additional accelerometers was outside of the budget. In future studies, increasing the number of accelerometer recordings, would increase measurement precision, as variable recording lengths per session and intermittent gyroscope channel dropouts on select days restricted data continuity. Additionally, the accelerometer used in this study has a maximum measurement limit of ±16 g, which is consistent with the peak recorded values (16.35 g). This hardware boundary means the sensor artificially truncated our absolute peak acceleration and force measurements. Implementing these technical improvements in future studies would provide increased precision for quantifying linear and rotational acceleration, as well as impact force measurements

Overall, every model has its limitations, but this preliminary study has determined that headbutting likely causes goats to begin the process of neurodegeneration at a young age, and it likely increases in severity over time. Common laboratory models can be ideal for studying the effects of certain variables, while themselves being invariable. But to understand the real-life repercussions of a variable and complex disease like TBI, it is beneficial to observe spontaneous occurrence in a real-life animal (Garner, 2014), such as our headbutting goats.

This study is the first important step towards developing a goat model of neurodegeneration. Our data suggests these animals sustain a lifelong accumulation of hundreds of thousands of repetitive mild head impacts with the potential to develop into chronic brain injury. We detected early neurodegeneration using clinically-relevant methods developed to target humans, indicating a high potential for translatability of the goat model, and a novel opportunity to understand the long-term consequences of repetitive TBI.

## CONCLUSION

This study investigated whether naturally occurring repetitive headbutting in goats is associated with behavioral, physiological, and histological markers of neurodegeneration. The detection and measurement of amyloid beta, phosphorylated tau, GFAP, and S100B in the CSF, alongside the corresponding immunohistochemical findings in brain tissue provides evidence that headbutting goats can accumulate the same biomarkers of neurodegeneration that are seen in humans and that this accumulation begins at an early age. While still in the early stages, the histopathological changes in the goat’s brain tissue are consistent with changes seen in human neurodegeneration caused by repetitive head impacts. Additionally, this is the first study of its kind demonstrating feasibility of PET/MRI in a goat TBI model, providing additional model characterization and opportunities for additional studies and clinical translation. Taken together, the biochemical and histological changes observed in our goats suggest the presence of early-stage neurodegenerative processes, likely reflective of a subclinical response to repetitive head impact. Despite the identified limitations, headbutting goats offer a unique and biologically relevant system for studying the early effects of repetitive, subclinical brain injury, and further study of this model has the potential to reduce the translational gap in neurodegeneration research.

## ACKNOWLEDGEMENTS

We are especially thankful to staff from UAB and MSU who assisted with the goat experiment, Domenica Pringle, Crystal Canady, Krista Pack, Randy Mills, and Aumbriel Schwirian, DVM, and the UAB cyclotron facility staff for PET tracer preparation. At UA, we thank Raghu Ganugula and his team as well as Stan Chtarbanova and Laura reed for providing experimental support. We thank Carmen Falcone for her insight on varicose projection astrocytes. We want to thank the following students from the CVN lab who performed manual counting of thousands of goat headbutts, listed in order of most headbutts counted: Emma Bowie, Sierra Lissick, Cameron Rancher, Claire Eddins, Alisa Kozyuk, Preston Herring, Rana Adam, James Blane, Brienna Ciemny, Maren McKean, and Connor Moody. Thank you to Rana Adam and Bre Ciemny for collecting accelerometer data. Finally, we thank the public supporters of our work, who donated to our lab during our headbutt-counting livestream: Jolene Paul, Reese Richardson, Eunice Campbell, Biocow, The Daily Tech News Show, Sunbun, PhillyCodeHound, and Jonathan Dombrosky. We would also like to acknowledge the late Lukasz Ciezla, a cherished friend and colleague who will be dearly missed.

