## supplemental information for "Headbutting goats self-inflict traumatic brain injury"

** Communicating author

**SUPPLEMENTAL METHODS**

The total resultants for linear acceleration (**a_mag_**), and rotational velocity kinematics (𝝎**_res_**) were calculated from the sensor data for each of the vectors (x-, y-, z-) (Equation 1 and 2).

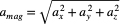
 ……… (Equation 1)

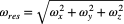
 ……… (Equation 2)

Force was calculated from a_mag_, where effective mass calculation for force estimation from assumes a standardized head-neck complex equals 8% of the subject's total body mass (Equations 3 and 4).

F = m_eff_ x a_mag_ ……… (Equation 3)

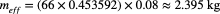
 ……… (Equation 4)

Total session impact impulse (**I**) represents the integrated area under the estimated force-time curve across the recording timeline. To isolate collision kinetic profiles from baseline movement, the summation loop was restricted strictly to active threshold rows where a_mag_ > Mean} + 2SD (Equation 5).

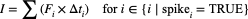
 ……… (Equation 5)

**SUPPLEMENTAL FIGURES**

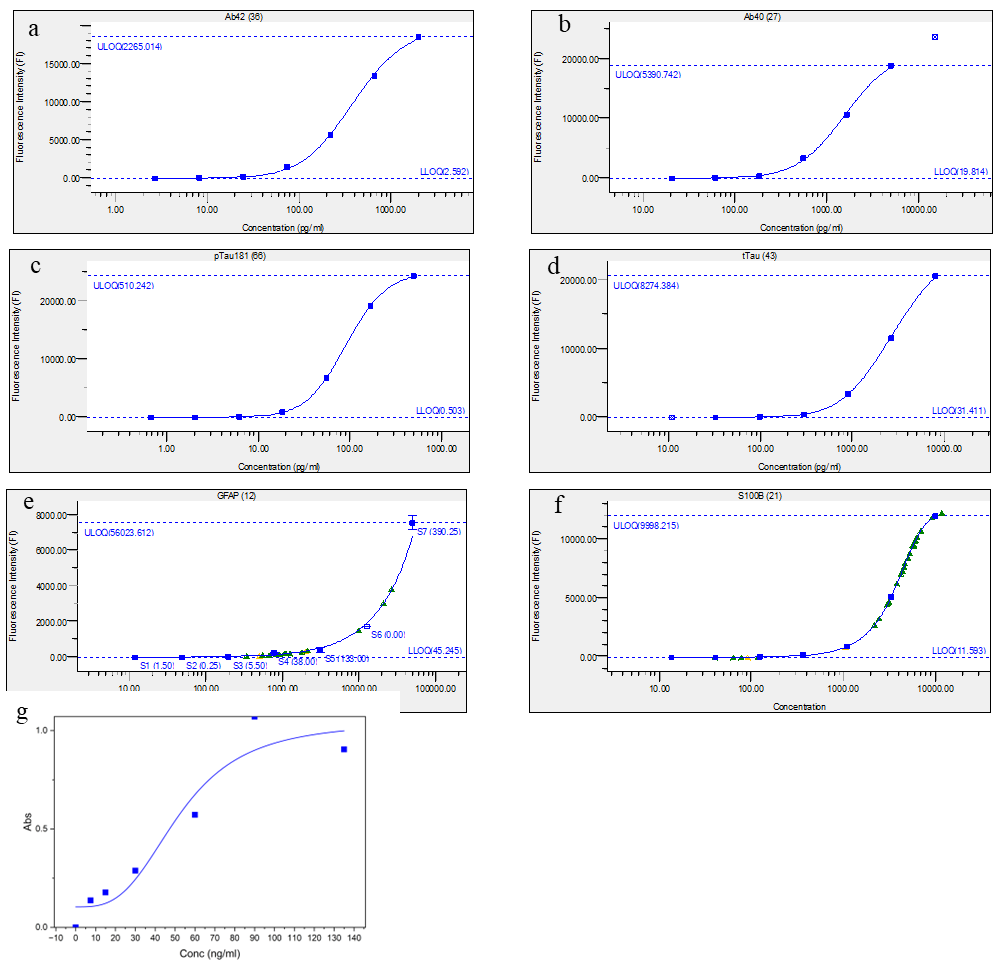

**Figure S1: Standard curves for biomarker assays.** Aβ_42_ (a), Aβ_40_ (b), pTau (c), tTau (d), GFAP (e), S100B (f) were measured with Multiplex bioassay while and NF-M (g) was measured by traditional ELISA.

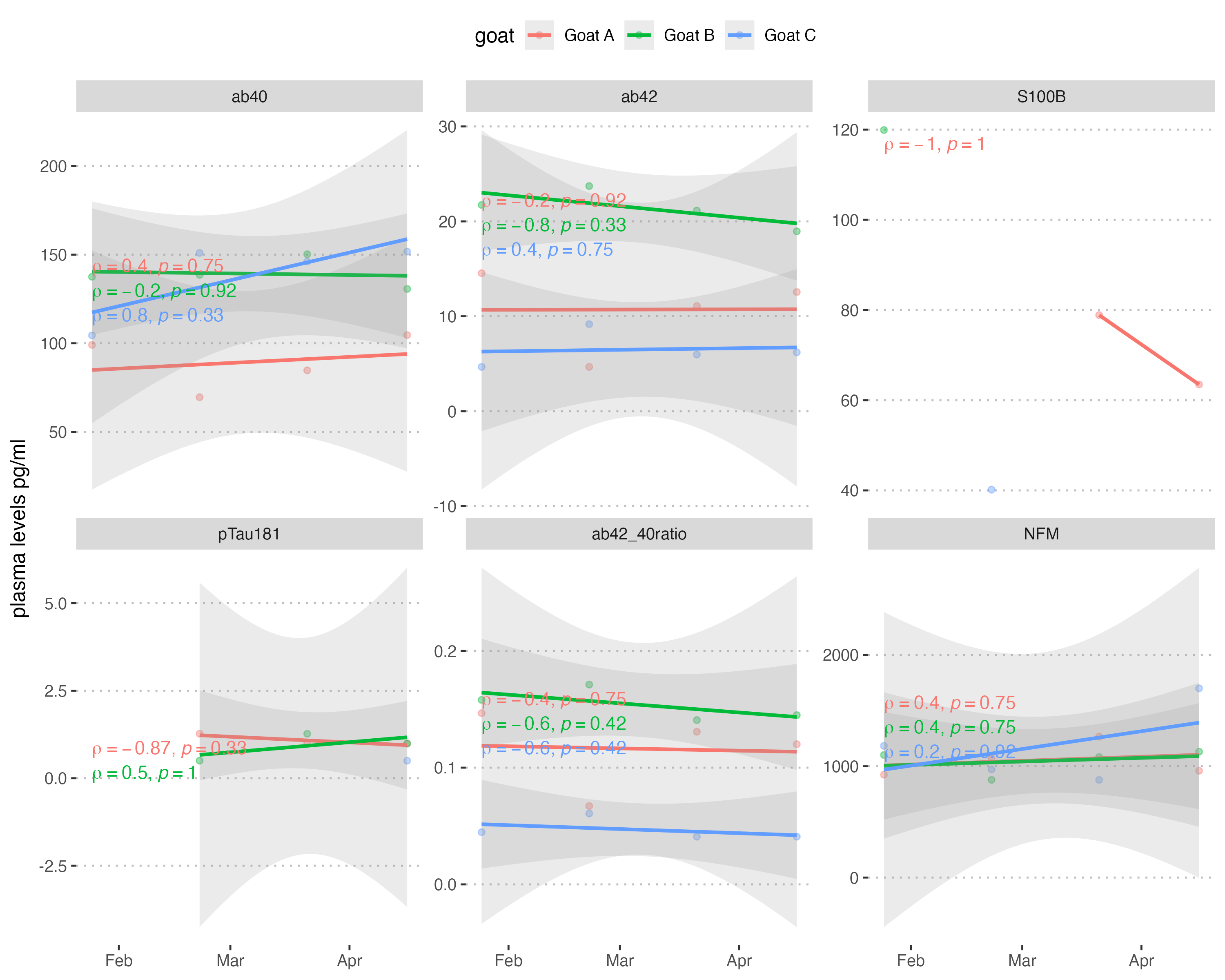

**Figure S2: Monthly plasma concentrations of biomarkers in headbutting goats (*Capra hircus*, n = 3)**. Biomarkers were measured from CSF samples using multiplex ELISA assays and regression lines show the longitudinal trends of the measured biomarkers.

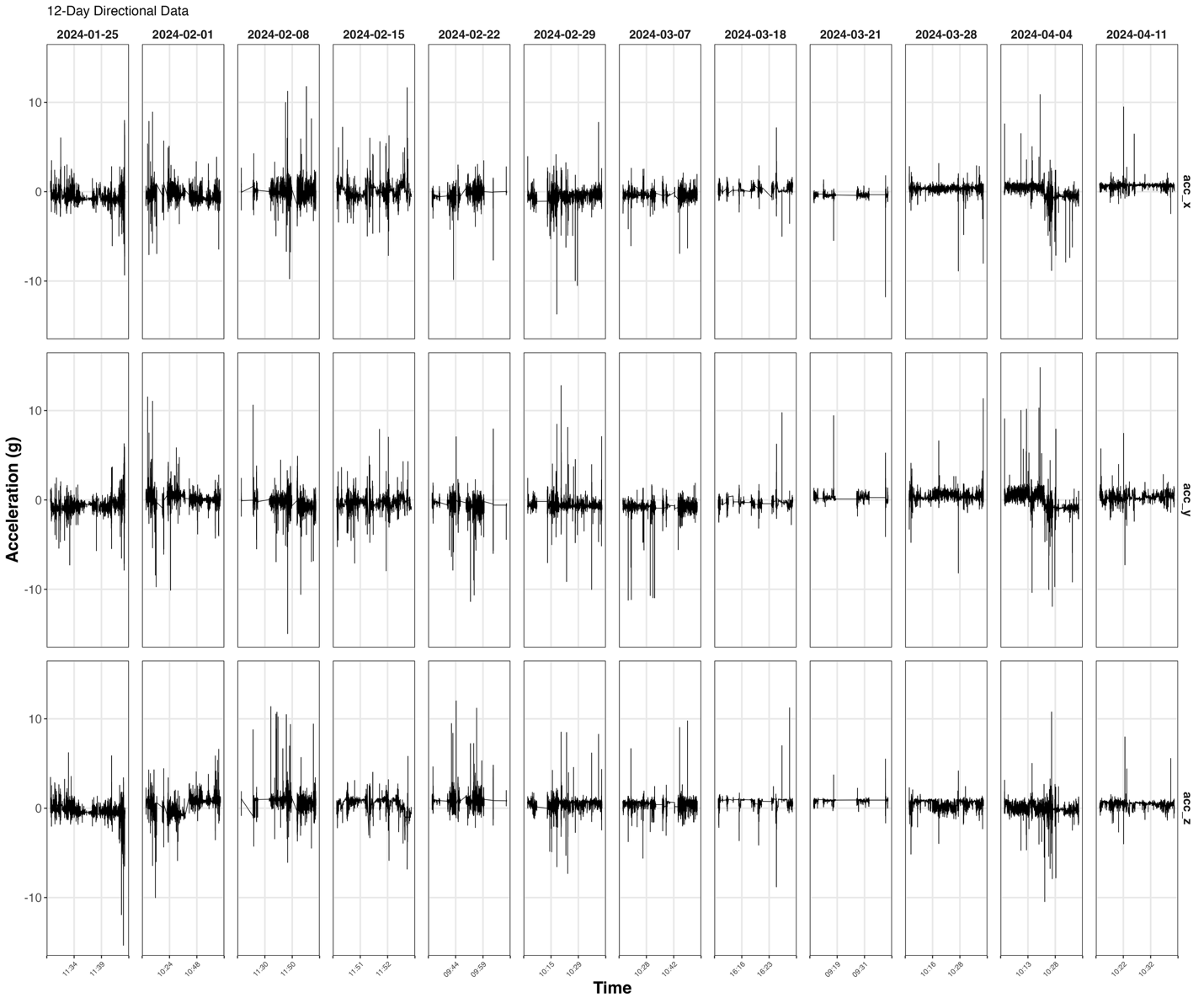

**Figure S3: 12 day acceleration data for goat A.** Directional acceleration measurements (in g) along the x- (acc_x), y- (acc_y), and z- (acc_z) axes.

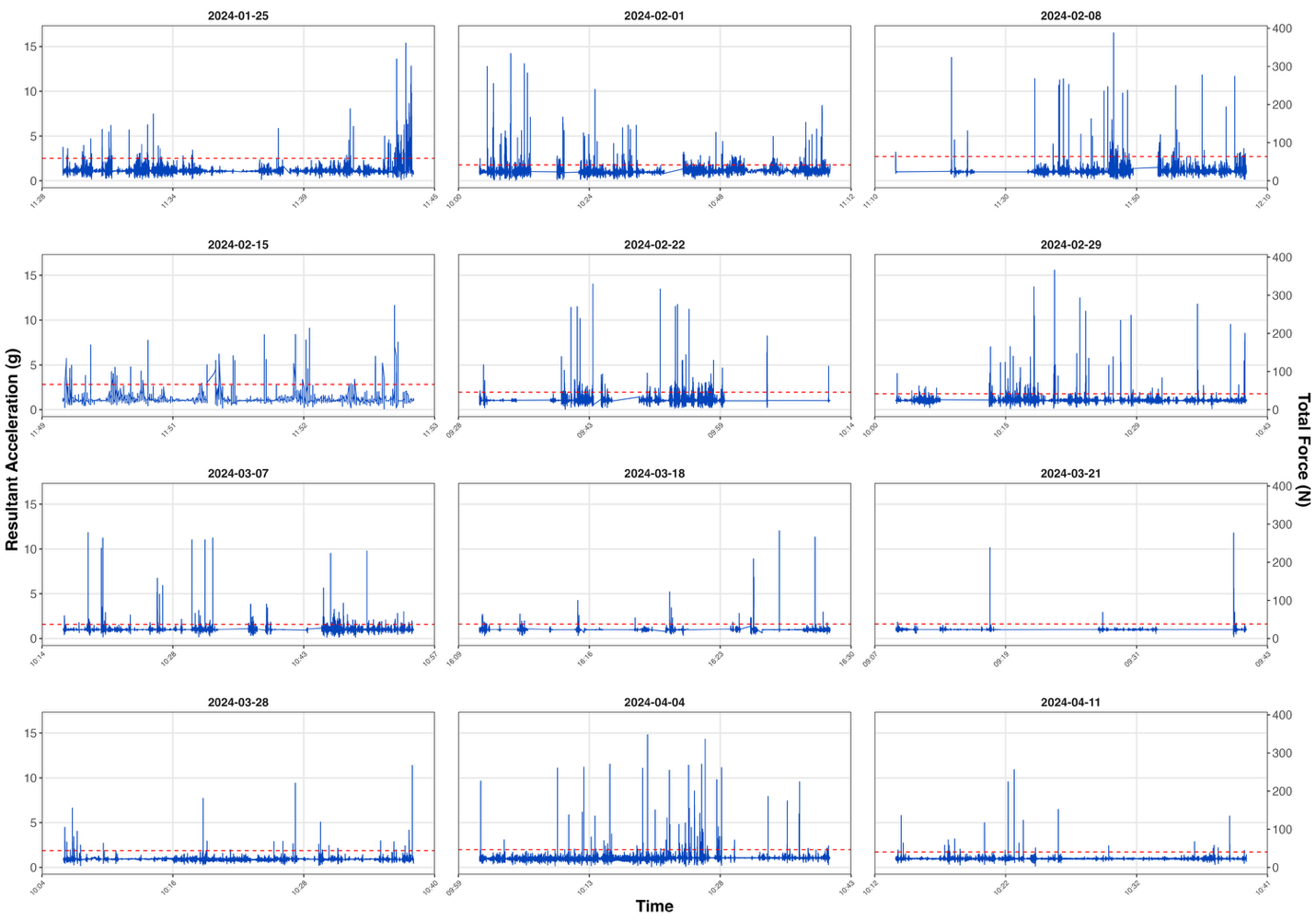

**Figure S4: 12 day resultant acceleration and estimated impact forces.** Longitudinal dual-axis arrays display the total resultant acceleration magnitudes (in g force) and linear impact forces (in N) across all 12 days, with horizontal dashed lines (in red) representing high-magnitude impact thresholds.

**Table S1: Biomechanical and Kinematic Metrics.** Summary of metrics calculated from accelerometer data for all 12 days in goat A. Note that angular velocity values with 0.00 denote time stamps with intermittent gyroscope channel dropouts, while linear accelerometer readings remained continuously active.

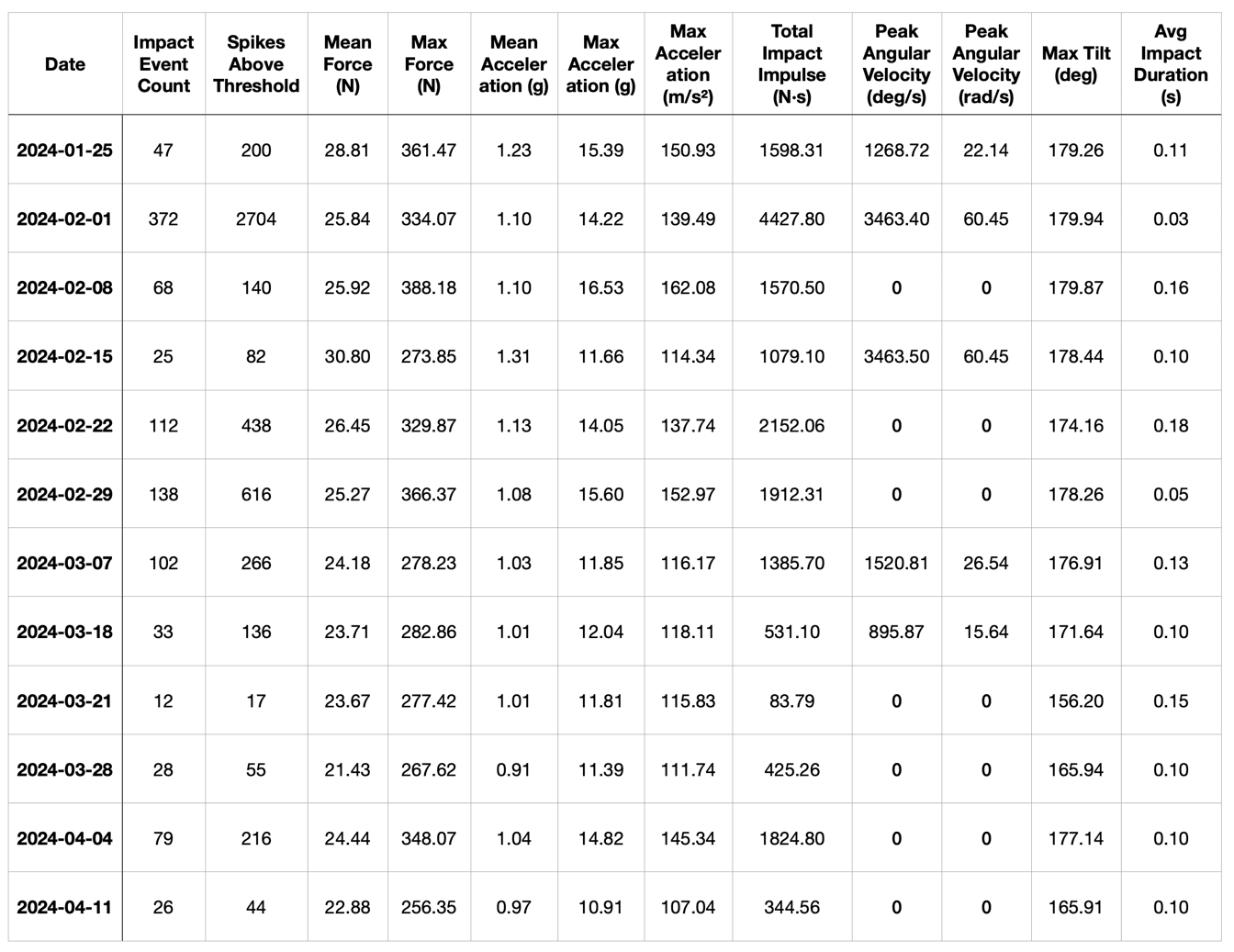
